# Host adaptation drives genome evolution and virulence diversification in a bacterial zoonotic pathogen

**DOI:** 10.64898/2026.06.29.734755

**Authors:** Alexandre Giraud-Gatineau, Kenneth Weke, Rania Ouazahrou, Francisco Pulido, Marc Monot, Nadia Benaroudj, Frederic Veyrier, Helena Pětrošová, Mathieu Picardeau

## Abstract

Understanding how zoonotic pathogens diversify across reservoir hosts remains a central question in evolutionary biology and infectious disease research. Here, we address this challenge using *Leptospira interrogans*, a globally distributed bacterial pathogen with an extremely wide range of animal reservoirs, as a model. After defining 13 distinct genogroups that largely align with serogroups, we selected the strongly host-adapted rodent-associated lineage, which is more frequently associated with fatal outcomes in patients, and the cattle-associated lineage for further analysis. A comprehensive approach integrating multi-omics analyses and host-specific infection assays showed that these two genogroups have followed distinct evolutionary trajectories associated with host specialization. Genomics demonstrated that specialized genogroups are genetically less diverse, characterized by divergence in membrane and signaling genes, and by the acquisition of host-adaptive functions. Changes in gene expression and protein production revealed distinct regulatory programs, predominantly affecting virulence pathways in rodent-borne lineage and stress responses in cattle-borne lineage. Consistently, rodent-borne lineage causes greater disruption of human epithelial barrier integrity and elicits an attenuated host-dependent macrophage inflammatory response relative to cattle-borne lineage. Collectively, these findings reveal distinct host-adaptive strategies and remarkable evolutionary plasticity in a major zoonotic bacterium, highlighting the central role of intraspecies heterogeneity in shaping host specialization.

## MAIN

Nearly three-quarters of emerging and re-emerging infectious diseases in recent decades have originated from animal reservoirs^1^, posing global health challenges with significant socioeconomic consequences^2^. Most studies have been largely shaped by viral systems, leaving bacterial zoonoses without an equivalent framework to explain how genomic and phenotypic variation drive host specialization and disease severity^3^. This gap is particularly evident in zoonotic bacteria that circulate across multiple reservoir hosts. Among them, *Leptospira interrogans* provides a powerful model for elucidating the mechanisms underlying host adaptation and zoonotic emergence.

Leptospirosis is one of the most widely distributed zoonotic bacterial diseases globally. The disease is considered re-emerging due to climate change, increasingly frequent flooding events and rapid urbanization, which collectively increase human exposure to reservoir species, particularly rodents in densely population areas^4,5^. The disease burden is especially high in tropical regions, where leptospirosis ranks among the leading bacterial zoonoses responsible for non-malarial febrile illness^6^. Its incidence is also rising in temperate regions, including Europe^7,8^. Although probably underestimated, leptospirosis is responsible globally for more than one million severe cases and 60,000 fatalities per year^4^ and inflicts substantial economic losses on livestock and dairy industries^9^.

Pathogenic *Leptospira* species infect over 160 mammalian species, including domestic, livestock, and wild animals^10,11^. Leptospires have also been detected in reptiles and other poikilothermic vertebrates such as snakes, frogs, toads and turtles^9^. This broad host range contributes to its persistence across diverse ecological niches^11^. Infected hosts are traditionally classified as natural (maintenance) or incidental. Maintenance hosts typically exhibit mild or no clinical symptoms but shed leptospires in their urine for extended periods, sometimes throughout their lifetime, thereby sustaining transmission cycle^12^. In contrast, humans, who are considered incidental hosts, may develop severe, sometimes fatal disease. However, these host categories are not always clear-cut, as intermediate states of host adaptation likely exist.

Host specialization is a defining feature of many zoonotic bacteria and is often shaped by long-term co-evolution between specific pathogen lineages and their reservoir hosts. In *Leptospira*, this co-adaptation has historically been associated with the surface-exposed O-antigen of lipopolysaccharide (LPS) which determines serovars. While more than 250 serovars have been described so far, only a minority have clearly identified reservoir hosts^10^. Thus, the serovar Icterohaemorrhagiae is associated with chronic and asymptomatic infections in the rat reservoir host, whereas this same serovar causes acute infections in other hosts^13,14^. However, as shared across other zoonotic bacterial pathogens, this serological classification alone does not capture the full extent of genomic diversity, regulatory variation and functional differentiation that underlie host specialization, immune evasion and disease severity^15,16^.

Beyond serological classification, *Leptospira* diversity is structured on the genomic level. There are nearly 80 species currently described; eight of which are clearly defined as pathogenic^17^. Among these, *Leptospira interrogans* is the most frequently associated with human infection worldwide^18^. Within *L. interrogans*, the serogroup Icterohaemorrhagiae has been associated with rodents reservoirs and represents the most severe clinical outcomes of leptospirosis in humans, including fatal cases^19–26^, raising fundamental questions about its ecological and pathogenic success. Despite its epidemiological and clinical importance, the genomic, transcriptomics, proteomic and host-interaction features distinguishing this serogroup from other *L. interrogans* serogroups remain unexplored, as is the case for all other serogroups. Addressing this gap is essential to understanding how intra-species diversity impacts pathogenicity and host specialization. Notably, the well-defined rodent reservoir of Icterohaemorrhagiae strains provides a model to investigate the evolutionary processes that drive host adaptation in zoonotic bacteria.

Here, we investigated the adaptive evolution, lineage differentiation and regulatory diversity of major *L. interrogans* genogroups using a combination of population genomics and comparative transcriptomics. By integrating global multi-omics data with functional analyses of epithelial barrier disruption and macrophage immune responses, we identify the determinants of intraspecies heterogeneity and host specialization. This combined approach provides broader insights into the evolutionary forces shaping persistence, virulence and host adaptation in one of the most globally distributed zoonotic pathogens.

## RESULTS

### Population structure and genogroup specialization in *L. interrogans*

To resolve the global population structure of *L. interrogans*, we conducted a phylogenomic analysis based on core genome single-nucleotide polymorphisms (cgSNPs) across 368 isolates (**Fig. 1a**). This dataset represents the most comprehensive collection to date, including strains from 12 serogroups, 25 diverse mammalian hosts in 45 countries and 5 continents (**Supplementary Fig. 1, Supplementary Table 1**). To ensure phylogenetic accuracy, we excluded intergenic recombination events and hypervariable regions such as the *rfb* locus encoding the O-antigen that determines serogroup and serovar^27^. This approach enabled us to focus on the conserved genomic signals driving evolutionary divergence.

**Figure 1.**
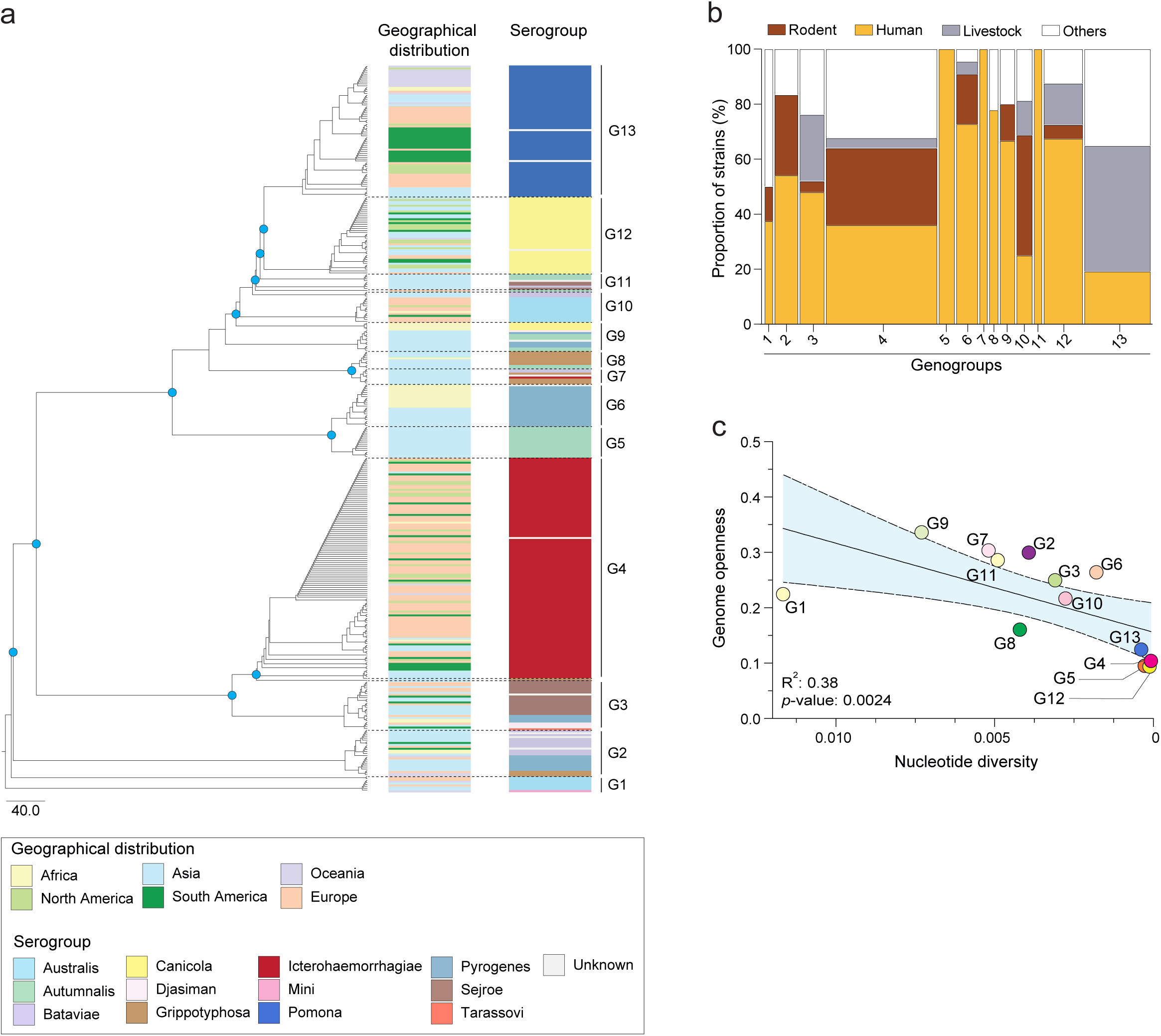
Population structure and ecological genome constraint of *L. interrogans* isolates. (a) Maximum-likelihood phylogeny of 368 *L. interrogans* genomes based on 5,666 core-genome SNPs (cgSNPs). The cgSNPs alignment was constructed after detecting and removing recombination events. The cgSNPs-based phylogeny tree is divided into 13 genogroups (G1-G13) delineated by clue node markers (bootstrap support >90%). Colored rings indicate serogroup assignments. The tree is rooted using *L. kirschneri* as an outgroup. (b) Distribution of host species distribution by genogroup. (c) Correlation between pangenome openness and nucleotide diversity. Two-sided Sperman’s rank correlation test was used, where the *p*-value, R^2^ (percentage of variance explained by the linear regression model) and the line and the shaded area depict the 95% confidence interval of the linear regression are represented in the panel.

This analysis revealed 13 distinct phylogenetic clusters, hereafter referred to as genogroups. Most genogroups closely matched the serological classification, indicating co-evolution between surface-exposed LPS and the core genome (**Fig. 1a**, **Supplementary Fig. 2**). For instance, genogroup 4 (G4) comprised isolates from the serogroup Icterohaemorrhagiae, while G5 and G13 aligned closely with serogroups Autumnalis and Pomona, respectively. Notably, some serogroups were distributed across multiple genogroups while maintaining their serovar identity. For example, isolates associated with serogroup Pyrogenes were distributed across G2, G3 and G6, with G2 containing the Manilae serovar and G3/G6 corresponding to the Pyrogenes serovar. These patterns suggest that genomic variation beyond the *rfb* locus contributes to the population structure and diversification of *L. interrogans*. Whereas most genogroups showed global distribution and broad host ranges, some displayed more localized geographic patterns and host restriction. For instance, G5 was exclusively identified in human cases in Laos, whereas G6, G7, and G11 were predominantly associated with human infections and confined to Asia (**Fig. 1a-b**, **Supplementary Fig. 1**).

Among all genogroups, G4 emerged as the most prevalent in our dataset, accounting for 30% (112/368) of the isolates, followed by G13 (18%; 67/368). The dominance of G4 is consistent with its frequent isolation in human severe cases of leptospirosis and its worldwide distribution in rats^10^ (**Fig. 1a-b**, **Supplementary Fig. 1**). Genogroup G13 exhibited a similarly broad geographic distribution but was more frequently associated with livestock, including bovine and pig reservoirs^9^.

We hypothesized that genogroups restricted to specific hosts or geographic regions exhibit reduced nucleotide diversity and limited gene acquisition, reflecting niche refinement^28^. Accordingly, analysis of coding sequences revealed important variation in nucleotide diversity across genogroups. Genogroups G1 and G9 exhibited high diversity, whereas G4, G5, G12 and G13 showed reduced diversity (**Fig. 1c**, **Supplementary Fig. 3a**). Genogroups with low diversity also had lower pangenome openness, with 4,015 core and accessory genes in G4,

G5, G12 and G13 compared to 4,913 in the more variable genogroups (**Supplementary Fig. 3b-c**). A correlation between pangenome openness and nucleotide diversity was observed, suggesting that G4, G5, G12 and G13 exhibit hallmarks of an ecological specialization (Spearman’s ρ= 0.76; *p*-value= 0.0024; **Fig. 1c**). Within this subset, G4 was predominantly isolated from rodents (51% of animal reservoirs). Apart from rare exceptions, this genogroup was not found in livestock (**Fig. 1b**, **Supplementary Table 1**). In contrast, G13 was largely associates with livestock (56% of animal reservoirs) and was not found in rodents. Collectively, these patterns provide evidence for host specialization in G4 and G13 genogroups, which also represent the most prevalent genogroups in our dataset. Accordingly, subsequent analyses focus on G4 and G13, hereafter referred to as G4-Ict and G13-Pom, respectively, in reference to their serogroup classification as serogroups Icterohaemorrhagiae and Pomona.

### Functional domain differentiation of specialized genogroups

To assess whether ecological specialization among genogroups is reflected in their functional protein repertoires, we analyzed protein domain composition across all isolates using a HMMER-based search. Principal component analysis (PCA) of protein domains revealed distinct genogroup-specific clustering for G4-Ict and G13-Pom (**Fig. 2a**). The domains contributing to the separation of genogroups were associated with phage-related elements, mobile genetic elements, metabolism and membrane remodeling (**Supplementary Fig. 4**, **Supplementary Table 2**). Notably, phage-associated and mobile genetic element domains were major contributors to functional differentiation across genogroups with high genetic diversity (G1, G2, G6, G7, G8, G9, G11). However, these domains were markedly reduced in G4-Ict and G13-Pom, which was consistent with reduced horizontal gene transfer capability and increased genomic stabilization in these host-specialized genogroups.

**Figure 2.**
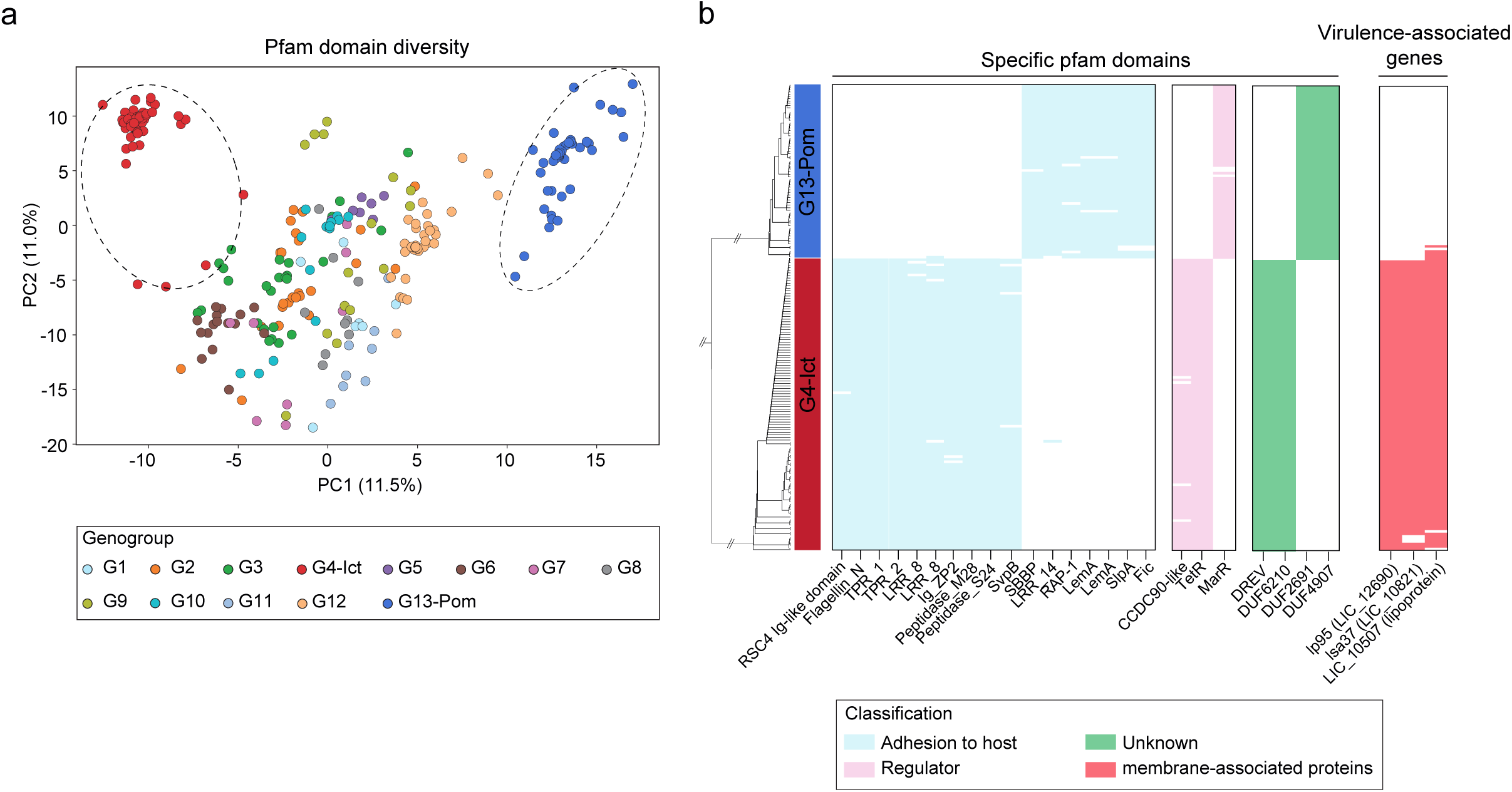
Protein domain variability in *L. interrogans*. (a) PCA plot of *L. interrogans* genomes based on Pfam domain composition. Each point represents a genome, colored by genogroup as indicated in the legend. Genogroups G4-Ict and G13-Pom are encircled to highlight genogroup-specific clustering. (b) Heatmap showing the distribution of specific Pfam domains in *L. interrogans* genogroup G4-Ict and G13-Pom, aligned with the cgSNPs tree. Categories significantly enriched or depleted in specific domain families are colored in blue (Motility, invasion, host adhesion), pink (Transcriptional regulator), green (domain unknown) and red (Virulence-associated genes).

To pinpoint the genetic basis of host specialization, we compared the orthologous genes of G4-Ict and G13-Pom isolates. The analysis recovered a shared core of 3,050 genes, as well as 219 genes unique to G4-Ict (63.5% assigned as hypothetical proteins, followed by 7.8% belonging to cell wall/membrane) and 131 unique to G13-Pom (79.4% assigned as hypothetical proteins, followed by 5% belonging to cell wall/membrane). Thus, the divergence between G4-Ict and G13-Pom was driven by a relatively small subset of genes, rather than a considerable modification of the genome. Importantly, protein domains distinguishing G4-Ict from G13-Pom were associated with functions potentially linked to host adaptation, including invasion, host adhesion and transcriptional regulation (**Fig. 2b**). For example, G4-Ict specific domains include peptidase families (Peptidase_M28 and Peptidase_S24) previously implicated in host invasion in *Leptospira*^29^, as well as leucin rich repeat (LRR_8), a common domain family for host proteins interaction^30^. G4-Ict also encoded an SpvB domain (LIC_12045) which is found in actin-ADP-ribosylating toxins^31^, a TetR-family transcriptional regulator (LIC_11584) potentially involved in host adaptation^32^ and a CCDC90-like domain (LIC_10110) a eukaryote-associated regulator of the mitochondrial calcium uniporter^33^. Similarly, G13-Pom was characterized by domains associated with structural and regulatory remodeling. Examples include the protein containing SlpA domain (LNFBFJNK_01030; see **Source Data Fig. 2** for gene nomenclature), which encodes a surface layer protein, essential for virulence in other pathogenic bacteria^34^ and protein containing the regulator MarR domain (LNFBFJNK_01305) that could be required for responses to environmental changes, such as those during host invasion^35^ (**Fig. 2b**, **Supplementary Fig. 5**).

In addition, analysis of virulence factors distribution revealed that while *L. interrogans* virulence genes, such as the lipoprotein *ligB*, the catalase *katE* and the collagenase *colA*^36^ were conserved across genogroups, some others virulence genes showed differential distribution in genogroups (**Fig. 2b, Supplementary Fig. 4**). Accessory genes involved in fibrinolytic modulation (*lsa37, LIC_10821*)^37^ and extracellular matrix adhesion (*lp95, LIC_10507*)^38^ were absent in G13-Pom compared to G4-Ict, suggesting genogroup-specific host interaction strategies. Together, these results show that specialized *L. interrogans* G4-Ict and G13-Pom encode distinct protein domain and virulence gene repertoires, potentially required in adaptation to specific ecological or host niches.

### Genogroup-specific variation in surface and signaling proteins drives host tropism

Building on this evidence of a niche-specific adaptation, we hypothesized that recurrent patterns of genetic variation across *L. interrogans* genogroups reveal functional signatures of evolutionary specialization, particularly among surface-exposed proteins that mediate host-pathogen interactions. Genome-wide analysis of single-nucleotide polymorphism (SNP) revealed heterogeneity of genetic variation in 479 genes constituting hypervariable “hotspots” (>60 SNPs; **Supplementary Fig. 6a**), which are broadly distributed along the genome (**Supplementary Fig. 7),** whereas 958 genes remained highly conserved (<10 SNPs). Cluster of orthologous gene (COG) enrichment of the highly polymorphic genes highlighted two categories that were significantly over-represented across all genogroups: Cell wall/membrane (48 genes) and Signal transduction mechanisms (43 genes; **Supplementary Fig. 6b-c**). These same functional categories were also significantly enriched within the host-specialized G4-Ict and G13-Pom (**Fig. 3a**), implicating surface remodeling and environmental sensing as primary targets of adaptive divergence. These SNP-enriched genes encoded lipoproteins, membrane proteins, histidine kinases and key virulence factors such as the leptospiral outer membrane OmpL1 and the surface adhesins LigB, LenA, Lsa77 and Lsa46, all implicated in host-pathogen interactions or immune evasion^36^ (**Fig. 3a**, **Supplementary Fig. 6c**, **Supplementary Fig. 8**). Elevated SNP densities were also observed in peptidoglycan synthesis and lipid A biosynthesis pathways, suggesting modulation of surface architecture (**Supplementary Fig. 6c**). Conversely, genes highly conserved across genogroups were enriched in essential cellular functions, including transcription (36 genes), energy production (39 genes) and coenzyme metabolism (39 genes; **Supplementary Fig. 9**), reflecting stringent selective pressure on core physiology.

**Figure 3.**
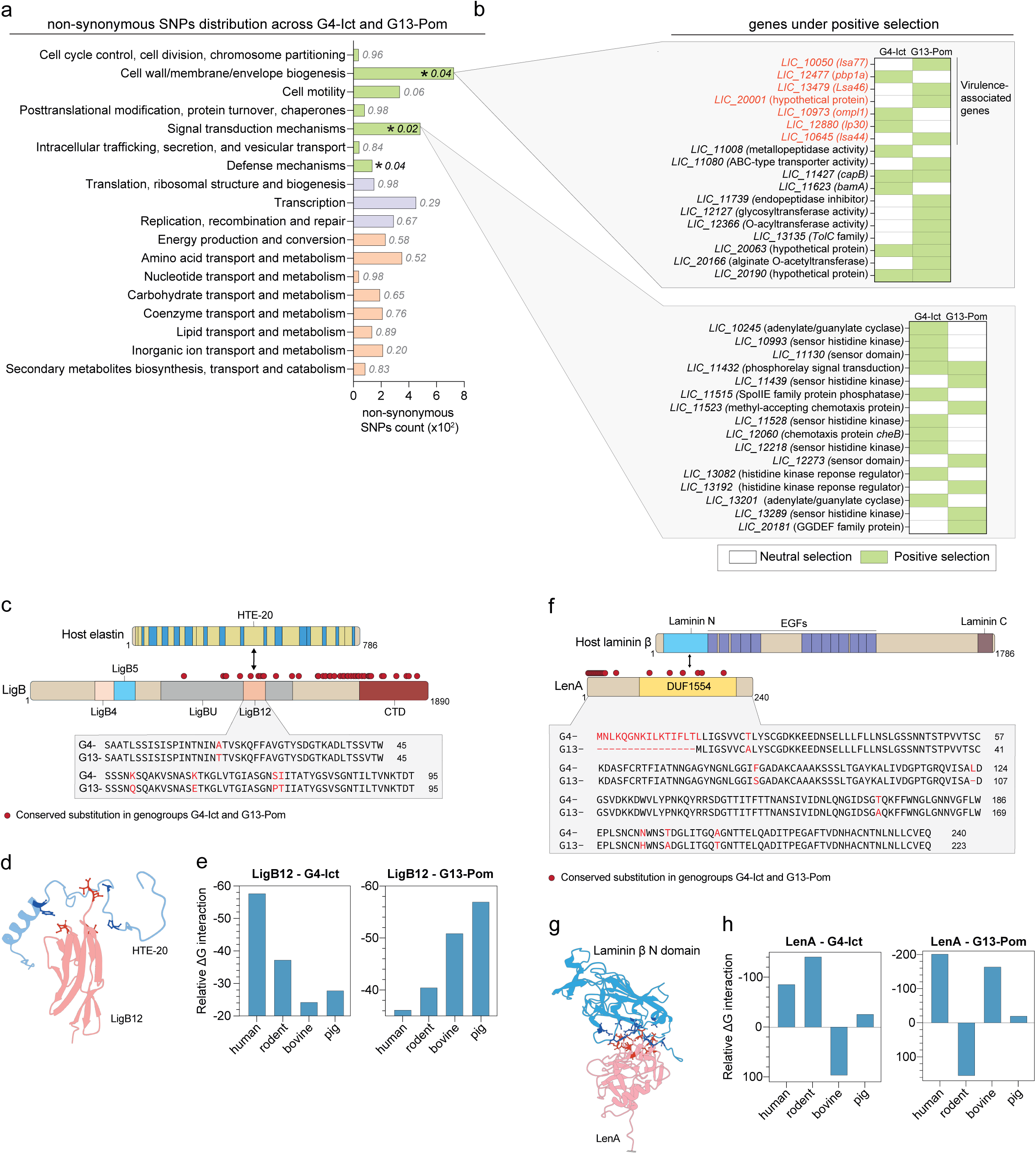
Adaptive genomic signatures in *L. interrogans* specialized genogroups G4-Ict and G13-Pom. (a) Functional categorization of genes enriched in non-synonymous SNPs in G4-Ict and G13-Pom using COG classification (green: Cellular process and signaling, purple: Information storage and processing, orange: Metabolism). Sub-categories are detailed on the y-axis. Fisher’s exact test with Benjamini-Hochberg method correction was used to estimate the enrichment of categories. *, *p* ≤0.05; **, *p* ≤0.01. (b) Non-synonymous SNPs-enriched genes under positive selection in G4-Ict and G13-Pom. Genes under positive selection belonging to Cell wall/membrane and Signal transduction mechanisms are indicated in green. Virulence-associated genes were labelled in red. (c) Domain-architecture representation of human elastin and *L. interrogans* LigB. The elastin schematic illustrates exon-derived repat modules (THE). Domains of the LigB protein are represented, with red dots indicating residue substitutions in genogroups G4-Ict and G13-Pom. The amino-acid sequences of the LigB12 domain from G4-Ict and G13-Pom are aligned and residues that differ are highlighted in red. Protein length in amino acid is indicated for each protein. (d) Model of the human elastin HTE-20 domain (in blue) in complex with the *L. interrogans* LigB12 domain from genogroup G4-Ict (in red). The two domains are displayed as cartoon ribbons. The interface residues identified as energetically important by FoldX analysis are highlighted in stick representation. (e) Predicted relative binding free energies (ΔG interaction) between the LigB12 from genogroup G4-Ict (left panel) or genogroup G13-Pom (right panel) and elastin HTE-20 domain from different hosts. Relative ΔG values were obtained using Rosetta InterfaceAnalyzer. Lower ΔG values indicated stronger predicted interactions. (f) Domain-architecture representation of human laminin β and *L. interrogans* LenA. Domains of both proteins are represented, with red dots indicating residue substitutions in LenA from G4-Ict and G13-Pom. The amino-acid sequences of LenA from G4-Ict and G13-Pom are aligned and residues that differ are highlighted in red. Protein length in amino acid is indicted for each protein. (g) Model of the human laminin β N-domain (in blue) in complex with *L. interrogans* LenA from genogroup G4-Ict (in red). The two domains are displayed as cartoon ribbons. The interface residues identified as energetically important are highlighted in stick representation. (h) Predicted binding free energies (ΔG interaction) between the LenA from genogroup G4-Ict (left panel) or genogroup G13-Pom (right panel) and the laminin β N-domain from different hosts.

Additionally, we observed that surface and signaling proteins variation was conserved by genogroup identity. Within enriched categories, disruptive mutations (frameshift, indels and missense) were reduced within individual genogroups, when compared to species-wide diversity (**Supplementary Fig. 10a**), indicating genogroup-specific conservation of allelic variants. Phylogenies reconstructed from hypervariable protein sequences recapitulated the cgSNP topology (**Supplementary Fig. 10b**), confirming that diversification of membrane and signaling proteins drives genogroup differentiation. Consistent with adaptive evolution involved surface and signaling proteins, genome-wide screen dN/dS analysis identified positive selection on cell wall/membrane categories in 7 of 13 genogroups (**Supplementary Fig. 6d**, **Supplementary Tables 3-4**), with peptidoglycan and lipid A pathways emerging as recurrent targets. Notably, host-specialized G4-Ict and G13-Pom showed positive selection on specific virulence-associated surface proteins (Lsa77 and OmpL1), and several histidine kinases (**Fig. 3b**), linking genetic diversification to ecological specialization.

To investigate whether conserved substitutions in G4-Ict and G13-Pom could directly participate in host specificity, we investigated their effects on host-protein recognition using structure-based interaction modelling. Recent studies have demonstrated that AlphaFold can reliably predict protein-protein interactions, including those in *Leptospira*^39,40^. We therefore focused on two previously studied proteins involved in host-pathogen interactions in *Leptospira*. The surface adhesin LigB from G13-Pom, through its LigB12 domain, binds exon 20 of the human elastin (HTE-20)^41^ (**Fig. 3c-d**) and the adhesin LenA from G4-Ict interacts with the human laminin^42^ (**Fig. 3f-g).** Screening of laminin subunits and domains identified the Laminin-N domain of the β subunit as the most likely interaction partner of LenA, based on the highest-confidence structural predictions **(Supplementary Fig. 11**). Structural prediction and binding affinity interface stabilities (ΔG) suggested genogroup-specific interface stabilities, which were dependent on the host species. The G4-Ict LigB12 domain was predicted to have a more stable binding (lower ΔG) to rodent and human elastins (HTE-20) compared to bovine or pig elastins. In contrast, G13-Pom LigB12 was estimated to bind in a more stable manner to bovine and pig elastin (**Fig. 3e**). Similarly, the complex between G4-Ict LenA and rodent laminin β N-domain displayed the highest stability, while the G13-Pom LenA exhibited a preferential binding affinity to bovine laminin (**Fig. 3h**). These predictions suggest that genogroup-specific sequence variation in surface adhesins likely determines host-tropism, providing a link between the observed genomic diversification and ecological specialization.

### Specialized genogroups exhibit distinct regulatory programs

Beyond genomic constraint, differences in gene expression may also contribute to ecological specialization. To further investigate genogroup specialization, we compared the transcriptomic profiles of five strains from G4-Ict and four strains from G13-Pom under host-like conditions^43^ (**Supplementary Table 5**). These strains were chosen based on their low number of *in vitro* passages, comparable growth kinetics (**Supplementary Fig. 12**) and maximal diversity in terms of geographic and host origin (**Supplementary Table 5**). To assess global transcriptional differences in these two genogroups, we first examined core genes that were differentially regulated under host-like (EMEM at 37°C) compared to *in vitro* (EMJH at 30°C) conditions within each genogroup, and second, genes that were differentially deregulated between the two genogroups under *host*-like condition.

Comparative transcriptomic analysis under host-like compared to *in vitro* conditions revealed a conserved core response shared by both genogroups, including 816 commonly upregulated (62%) and 802 downregulated (63%) genes, with enrichment in porphyrin, fatty acid, and sphingolipid metabolism, pathways previously associated with *Leptospira* adaptation to host-related conditions^43^ (**Supplementary Fig. 13a-b**; **Supplementary Tables 6-7**). Interestingly, ∼60% of SNP-enriched genes also showed differential expression under host-like growth (**Supplementary Fig. 14a**), highlighting their potential relevance. Despite this shared response, genogroups exhibited distinct transcriptional patterns. G4-Ict uniquely upregulated 303 and downregulated 288 core genes, compared to G13-Pom (*p*.adj ≤0.05), with an enrichment in teichoic acid biosynthesis, type II secretion systems, fatty acid metabolism and biofilm formation (**Supplementary Fig. 13a, c**). Furthermore, G13-Pom showed 198 up- and 188 downregulated genes, with enrichment limited to downregulated pathways associates to chemotaxis, flagellar assembly and fatty acid metabolism (*p*.adj ≤0.05; **Supplementary Fig. 13a, d**). Most genogroup-specific deregulated genes encoded hypothetical proteins and transposases. However, several genes encoded functionally relevant proteins, including lipoproteins and genes containing previously identified domains, such as SpvB (*LIC_12045*) and the TetR regulator (*LIC_11584*) in G4-Ict, as well as the genes containing the SlyA (*LNFBFJNK_01030*) and MarR (*LNFBFJNK_01305*) domains in G13-Pom (**Supplementary Fig. 14b-d**).

While these analyses revealed genogroup-specific transcriptional responses to host-like conditions compared to *in vitro* growth conditions, they did not determine whether G4-Ict and G13-Pom differ intrinsically in their expression programs when exposed to the same environment. To address this question, we directly compared gene expression profiles of G4-Ict and G13-Pom strains grown in host-like condition. This inter-genogroup analysis identified 70 genes upregulated and 134 genes downregulated (*p*.adj ≤0.05; |Log_2_FC| >1) in G4-Ict compared to G13-Pom (**Fig. 4a**, **Supplementary Table 8**). Notably, several key virulence factors of *L. interrogans* were mainly upregulated in G4-Ict. For example, genes encoding membrane proteins LigB^44^ and MPL36^45^ and the two major virulence-modifying (VM) proteins LIC_12340 and LIC_12339^46^ were strongly induced (**Fig. 4b**, **Supplementary Table 8**), reflecting enhanced pathogenic potential of G4-Ict. Consistent with this pattern, the most upregulated genes of G4-Ict encoded surface-exposed lipoproteins (**Supplementary Fig. 15**), supporting a role in host interaction and colonization. In contrast, genes upregulated in G13-Pom were predominantly associated with protein quality control (*clpB* and *hsp15-like*) and metabolic remodeling, whereas canonical virulence-associates genes were largely absent compared to G4-Ict (**Supplementary Fig. 15**). Proteomic analysis in the host-like condition confirmed the trends observed by transcriptomics. Lipoproteins and virulence-associated factors were enriched in G4-Ict, while chaperones and factors of the SOS response were detected in increased abundance in G13-Pom (**Fig. 4c-d, Supplementary Table 9**). Collectively, these complementary analyses highlight two distinct host adaptation strategies in these genogroups. The first, employed by G4-Ict, promotes the expression of factors involved in host interaction and dissemination. The second, associated with G13-Pom, enhances bacterial persistence and recovery following exposure to stressful conditions.

**Figure 4.**
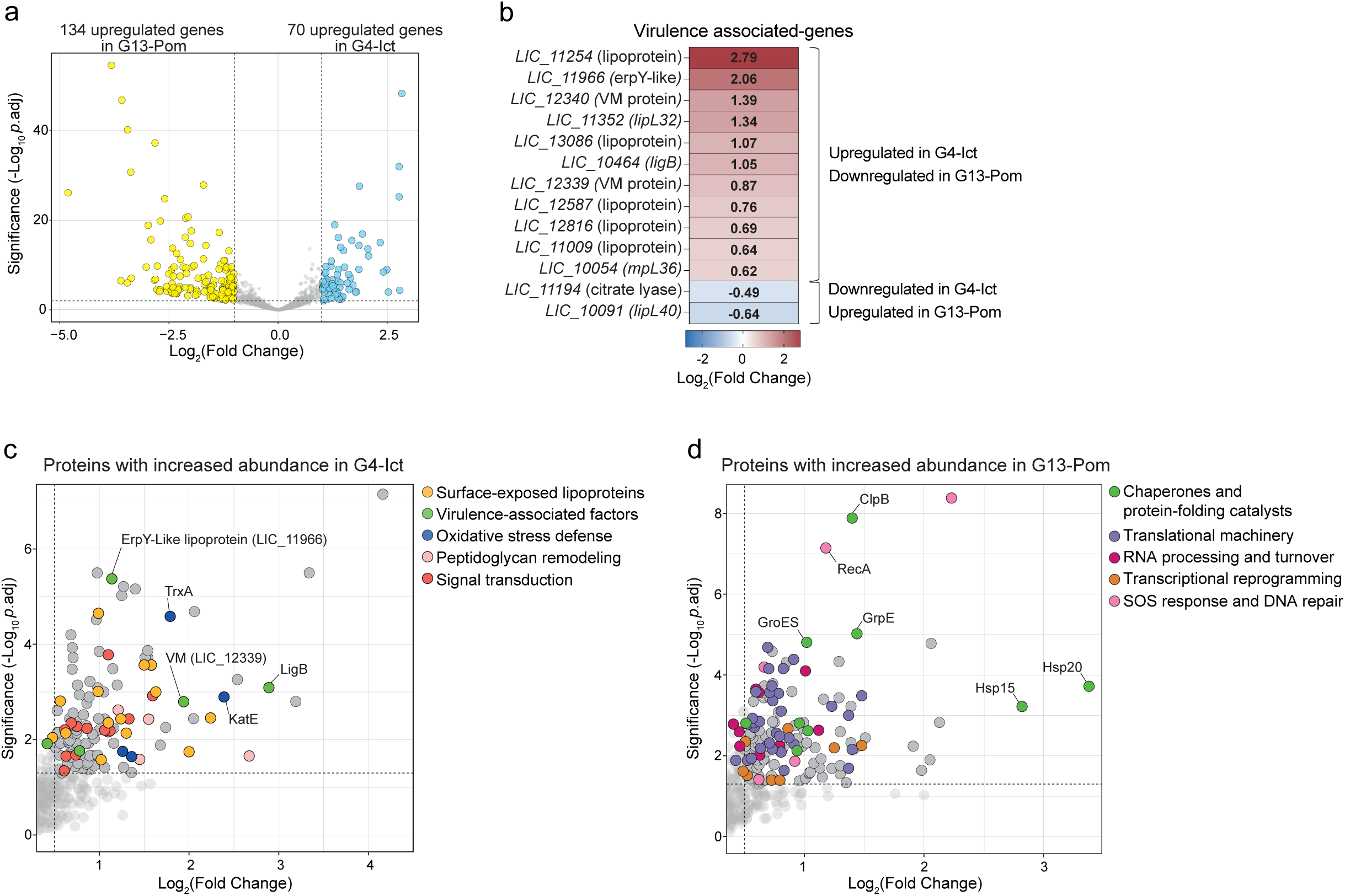
Comparative transcriptional and proteomic landscapes of *L. interrogans* genogroups G4-Ict and G13-Pom. (a) Volcano representation of *L. interrogans* DEGs of G4-Ict compared to G13-Pom in EMEM medium. Upregulated genes in G4-Ict and upregulated genes in G13-Pom are represented in blue and yellow, respectively, and genes that are not significantly differentially expressed are shown in grey (*p*.adj ≤0.05; |Log2FC| >1). (b) Heatmap of virulence-associated genes differentially expressed in *L. interrogans* G4-Ict compared to G13-Pom in EMEM medium (*p*.adj ≤0.05). Gene names are indicated on the left. Differential expressions are expressed as Log2FC with a color gradient from blue to red indicating low to high Log2FC. Value of Log2FC is indicated into the heatmap. (**c-d**) Volcano plots showing proteins significantly more abundant in G4-Ict compared with G13-Pom (c) and in G13-Pom compared with G4-Ict (d) in EMEM medium. The x-axis indicates the Log2 fold change in proteins abundance and the y-axis shows the -log10 adjusted *p* value (*p*.adj). Proteins with *p*.adj ≤0.05 are highlighted in darker colors. Dot colors represent distinct protein functional families as indicated in the legend of each volcano plot.

### Specialized *L. interrogans* genogroups differentially disrupt epithelial cell-cell junctions

Previous work has shown that the VM proteins LIC_12340 and LIC_12339 contribute to the disruption of adherens junctions^46^. Given the marked upregulation of these two genes in G4-Ict compared to G13-Pom, we next examined whether these specialized *L. interrogans* genogroups differ in their ability to compromise epithelial integrity. Immunofluorescence microscopy revealed that both genogroups reduced membrane-associated cadherin levels at 24 hr post-infection in human epithelial cells, indicative of junctional disruption (**Fig. 5**). However, G4-Ict strains induced a significantly greater loss of cadherin (1.8-fold more) compared to G13-Pom strains. These findings demonstrate that G4-Ict isolates possess an enhanced capacity to disrupt epithelial barriers relative to G13-Pom in human cells, which may underlie their higher prevalence in human infections. Thus, ecological specialization among genogroups is accompanied by distinct capacities to impair host epithelial integrity.

**Figure 5.**
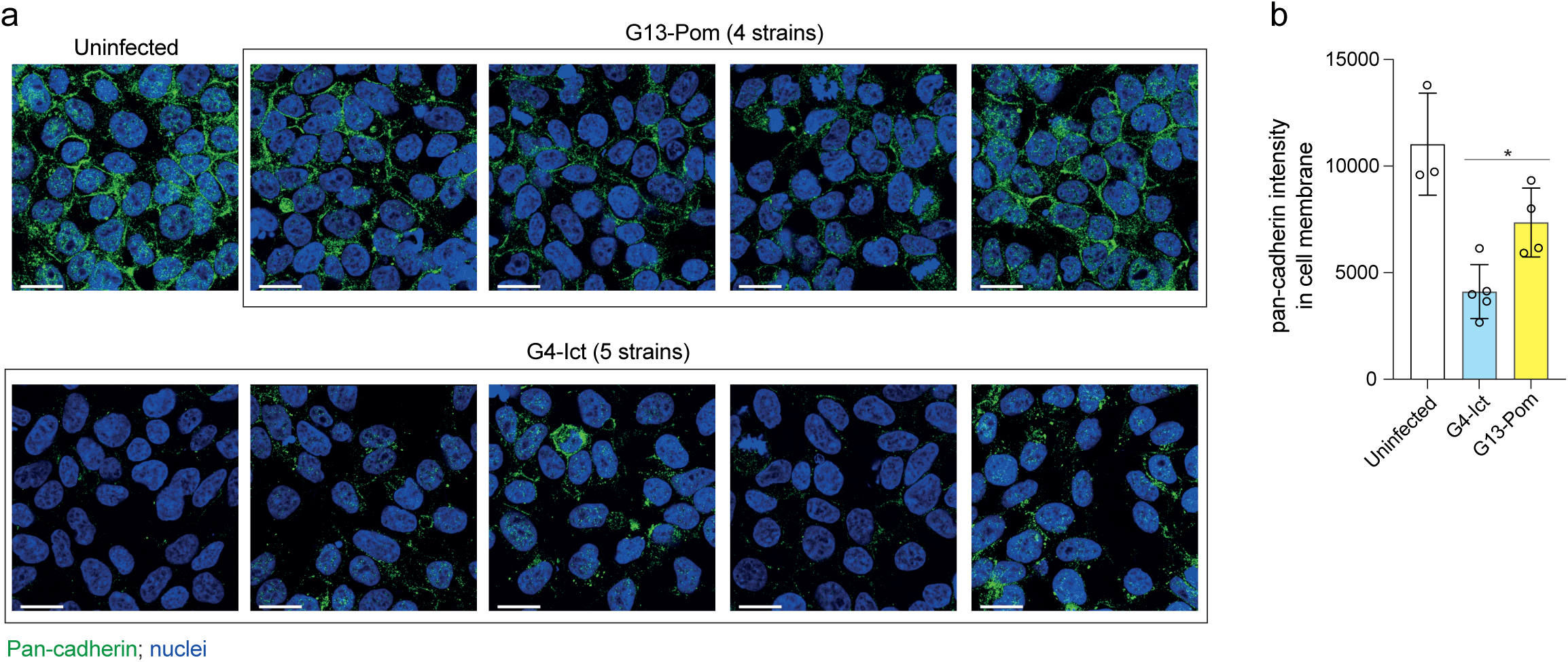
Specialized genogroups display distinct capacities to disrupt host epithelial barriers. (a) Confocal images of human epithelial cells uninfected or infected with *L. interrogans* strains from G4-Ict (5 strains) and G13-Pom (4 strains) for 24 hr (left panel), stained for pan-cadherin (in green) and DNA (in blue). Scale bars, 20 µm. The images are representative of 3 biological replicates. (b) Fluorescence quantification of pan-cadherin colocalizing with plasma membrane in epithelial cells uninfected and infected with G4-Ict or G13-Pom strains. Data represent the mean ± SD of 3 biological replicates. *, *p* <0.01 (unpaired, tailed *t* test).

### Specialized genogroups induce differential macrophage immune response

Next, we investigated whether these ecological specialized genogroups exhibit distinct immunomodulatory responses during host interactions by measuring comparative transcriptional response in mammalian macrophages. Infections of human macrophages revealed an amplified inflammatory transcriptional response with G13-Pom isolates compared to G4-Ict isolates as observed by the prominent induction of canonical inflammatory mediators, including *cxcl10*, *cxcl11*, *il6* and *il12b* (**Fig. 6a**; **Supplementary Table 10**). The increase expression of several inflammation-associated genes was confirmed by RT-qPCR, supporting the robustness of the transcriptomic analysis (**Supplementary Fig. 16**). Pathway enrichment analysis further confirmed that the immune responses were not limited to few isolated cytokines but reflected a broad deregulation of host immunity. Significant enrichment of interferon-γ and -α responses, TNF-α signaling via NF-κB, inflammatory response, IL-6-JAK-STAT3 signaling, complement and apoptosis signatures was detected in G13-Pom-infected cells, which reflected a coordinated activation of multiple innate immune pathways (**Fig. 6b**). Indeed, G13-Pom strains exhibited enhanced pathogen-sensing pathways, with upregulation of nucleic acid and danger sensors such as *ddx58*, *ifih1*, *myd88* and *nod2*. A robust type I interferon-associated program and a pronounced inflammasome/ IL-1 axis including *il1a*, *il1b*, *casp1*, *nlrp3* and *il1rn* were also upregulated (**Fig. 6c**). Moreover, this immune activation was associated with an induction of inflammatory effectors, notably *tnf*, *il6*, *cxcl1, cxcl2, 3, 5, 6* and *ccl2, 3, 4, 5, 20*, which was consistent with a program link to myeloid and neutrophil recruitment. Together, these data show that relative to G4-Ict, G13-Pom elicits a broader and stronger inflammatory response in human immune cells. These enhanced responses may promote more rapid control of infection with G13-Pom, resulting in reduced disease severity compared to G4-Ict. In contrast, G4-Ict appears to possess a greater capacity to evade host immune recognition, which may contribute to its increased disease severity. However, this difference in gene expression did not correlate with a difference in increased survival rates in the presence of human serum (**Supplementary Fig. 17**).

**Figure 6.**
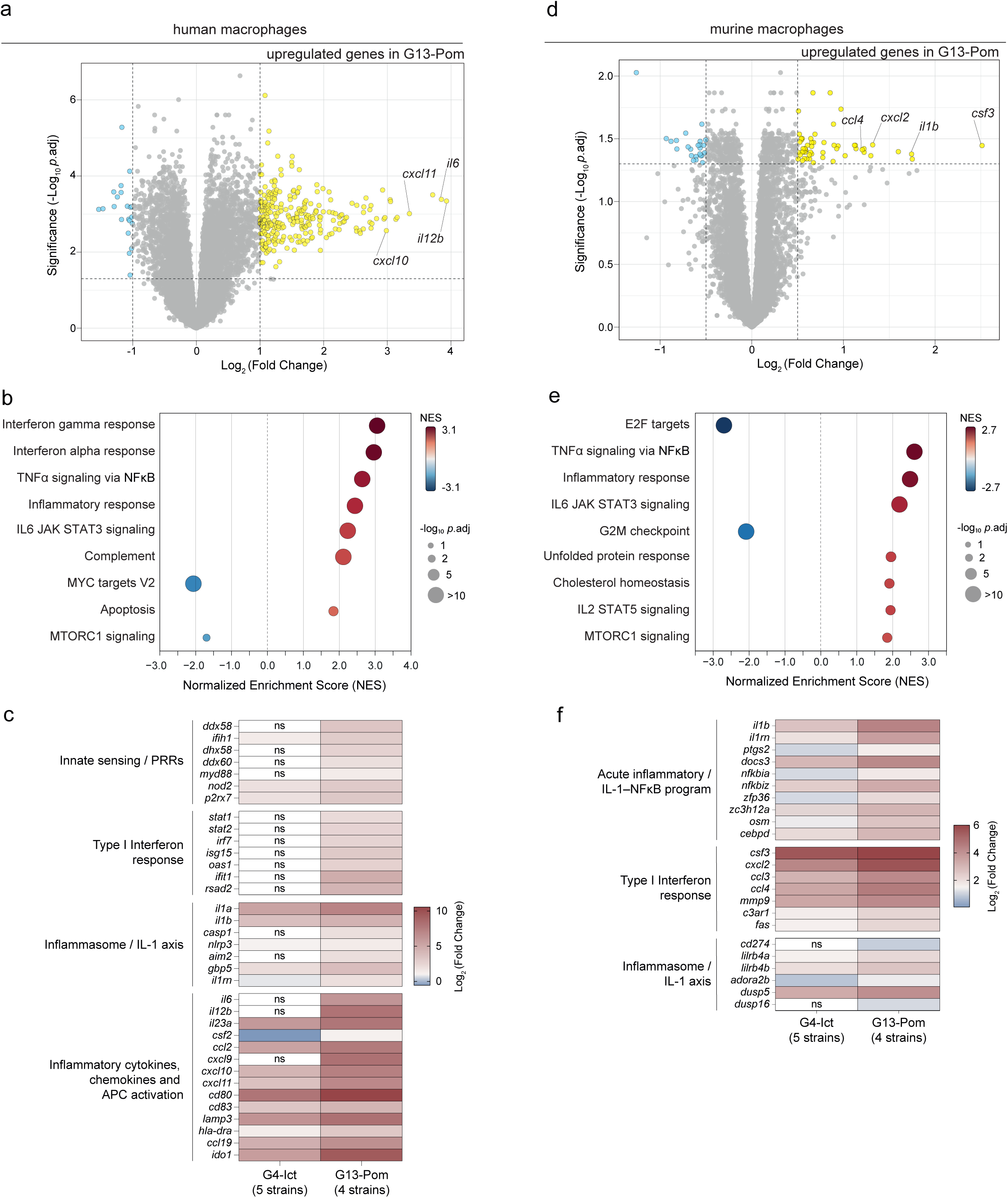
Genogroup-specific induction of innate immune responses in mammalian macrophages. RNA-Seq analysis in human and murine macrophages infected with *L. interrogans* strains from genogroups G4-Ict (n= 5 strains) and G13-Pom (n= 4 strains) at 6 hr post-infection. (a) Volcano representation of DEGs from G4-Ict-infected human macrophages compared to G13-Pom-infected human macrophages. G4-Ict and G13-Pom deregulated genes are represented in blue and yellow, respectively and not significantly DEGs are shown in gray (*p*. adj ≤0.05; |Log2FC| >1). Most deregulated genes associated are labelled. (b) Pathway enrichment analysis of G4-Ict-infected human macrophages compared to G13-Pom-infected human macrophages. The y-axis shows the enriched pathways, the x-axis values are the normalized enrichment score for each pathway, and the size of each circle represent the *p*.adj value. (c) Heatmap showing the most DEGs related to each category indicated in the right. Differential expressions are expressed with a color gradient from blue to red indicating low to high Log2FC. (d) Volcano representation of DEGs from G4-Ict-infected murine macrophages compared to G13-Pom-infected murine macrophages. G4-Ict and G13-Pom deregulated genes are represented in blue and yellow, respectively and not significantly DEGs are shown in gray (*p*. adj ≤0.05; Log2FC| >0.5). Most deregulated genes associated with inflammation are labelled. (e) Pathway enrichment analysis of G4-Ict-infected murine macrophages compared to G13-Pom-infected murine macrophages. The y-axis shows the enriched pathways, the x-axis values are the normalized enrichment score for each pathway, and the size of each circle represent the *p*.adj value. (f) Heatmap showing the most DEGs related to each category indicated in the right. Differential expressions are expressed with a color gradient from blue to red indicating low to high Log2FC.

To assess whether the differential immune response between genogroups was conserved across the G4-Ict reservoir host, we repeated the experiments using murine macrophages. Although the amplitude of inflammatory response between G4-Ict and G13-Pom was more restricted than in human macrophages, G13-Pom isolates still induced a higher inflammatory response in murine macrophages; expression of selected inflammation-associated genes was confirmed by RT-qPCR (**Fig. 6d**; **Supplementary Fig. 16**). We observed the upregulation of key inflammatory mediators including *csf3*, *il1b*, *cxcl2* and *ccl4* and a significant enrichment pathway related to TNF-α signaling via NF-κB, inflammatory response and IL-6-JAK-STAT3 signaling (**Fig. 6d-e**; **Supplementary Table 11**). Consistent with these enrichments, we observed an induction of genes related to IL-1/NF-κB pathways such as *il1a*, *il1rn*, *ptgs2* and *nfkbia*, as well as an induction of genes related to inflammasome/ IL-1 axis (**Fig. 6f**). Thus, as observed in human cells, G13-Pom promotes a more inflammatory state than G4-Ict in the mouse model, but this response is dominated by an IL-1 and innate stress-response program rather than broader strong inflammatory and antigen-presentation-associated profiles, as observed in human macrophages. Together, these results demonstrate that *L. interrogans* genogroups differ in their capacity to modulate host immune responses in a genogroup- and host-specific manner. Notably, lipid A profiling under host-like conditions revealed no structural differences between G4-Ict and G13-Pom, excluding a role for lipid A in these differential immunomodulatory phenotypes (**Supplementary Fig. 18**).

## DISCUSSION

Host specialization is central to the emergence of zoonotic pathogens, yet the evolutionary mechanisms shaping bacterial virulence and adaptation across diverse hosts remain poorly understood. Here, using *L. interrogans* as a model, our study provides insights into the evolutionary trajectories that drive bacterial zoonotic emergence. We show that host-associated diversification within a bacterial pathogen species emerges through the interplay of genomic constraints, adaptive changes in specific biological functions, and distinct regulatory programs. Together, these processes generate intraspecific heterogeneity, promoting specialization to different reservoir hosts and influencing severity to infection in humans.

Within the *Leptospira* genus, *L. interrogans* is the species most frequently associated with human infections worldwide and displays a wide diversity of serovars associated with multiple animal reservoirs. Here, using a collection of strains from different origins, we delineated 13 genogroups that capture both phylodynamic structure and ecological differentiation. Although these genogroups mostly correlate with serogroups, this classification reflects deeper phylogenetic link that goes beyond the O-antigen structure. While serotyping remains a powerful epidemiological tool, it does not fully account for lineage-specific differences in host adaptation or disease severity^47,48^. Notably, some genogroups sharing the same serogroup exhibited diverse geographic distributions, consistent with local adaptation to specific reservoir or environmental contexts. However, limited information exists on associations between *L. interrogans* isolates and disease severity, largely due to the challenges of strain isolation and inter-individual variability in susceptibility to infection. These constraints highlight the need for a comprehensive study integrating phylogenomic comparisons with *in vitro* and *in vivo* phenotyping studies.

Variation in pangenome openness across *L. interrogans* genogroups offers further insight into their lifestyles, as observed in other pathogens^28,49^. Genogroups with reduced pangenome openness and limited genetic diversity are associated with specific host or geographic niches, including rodent reservoirs (G4-Ict), human infections in Southeast Asia (G5 and G12), and livestock hosts (G13-Pom). The patterns observed suggest that these lineages have undergone ecological specialization, acquiring unique protein domains, with reduced genomic plasticity. The emblematic case is the Icterohaemorrhagiae serogroup, which is globally distributed, with rats as the primary reservoir, and whose isolates are highly genomically conserved^50^. The same scenario may apply to G13-Pom, which likely disseminated globally via pigs and cattle. In contrast, genogroups with more open pangenomes are likely shaped by fluctuating environmental pressures, requiring greater genomic flexibility for their persistence. In *Leptospira*, pathogenic species likely emerged during the evolution of mammals, allowing associations with specific hosts to shape bacterial genome dynamics over hundreds of millions years^14^. This divergence in genome dynamics thus suggests that prolonged evolutionary stability within a defined host-reservoir system may favor genome streamlining, whereas genogroups exposed to more variable ecological conditions maintain higher levels of genomic plasticity.

Adaptive evolution of zoonotic pathogens is tightly linked to host immune pressures and the need for efficient interaction with host tissues^51,52^. In addition to LPS, which plays a key role in host adaptation^13^, we identified hypervariable hotspot genes, with notable enrichment in membrane proteins and signal transduction. These genes are not only variable but also frequently under positive selection for specific genogroups, suggesting their important role for adaptation to a specific niche. These hotspot genes points to the possibility that long-term interaction with distinct host environments or reservoir species has driven the diversification of surface-exposed bacterial proteins. Computational modeling of LigB and LenA, two membrane-exposed proteins, reveals that conserved substitutions between genogroups drive distinct binding stabilities to host-specific extracellular matrix proteins elastin and laminin. Specifically, the predicted interface stabilities demonstrate the preferential binding of G4-Ict variants to rodent and human substrates, while G13-Pom variants exhibit preferential binding to bovine and porcine substrates. These findings suggest that sequence variations in surface-exposed proteins could be an additional molecular determinant of host tropism. Experimental characterization of the interaction between variants of proteins such as LigB and elastin or fibronectin from different mammalian species is needed to clarify whether these variations enhance host specificity or pathogenic potential.

Differences in virulence and host specificity may be explained not just from genomic content but from how genes are regulated. In pathogens, such as *Listeria monocytogenes*, lineage-specific regulation of stress responses and virulence can drive phenotypic heterogeneity without genomic differences^16^. We have previously shown that although the *katE* gene is present in both P1+ (high-virulence pathogens) and P1− (low-virulence pathogens) *Leptospira* species, it is differentially expressed between the two groups, which results in higher resistance to oxidative stress in P1+^53^. By sensing the host environment, pathogens adjust gene expression and metabolism to support survival and replication^53–55^. These regulatory strategies could reflect ecological adaptation and explain differences between zoonotic lineages. Consistent with this adaptation hypothesis, our transcriptomic and proteomic analysis of *L. interrogans* under host-like conditions revealed genogroup-specific regulatory patterns that are not detectable through genomic comparisons alone. G4-Ict showed upregulation of multiple host-adaptation pathways including type II secretion systems, lipoproteins and several key virulence factors. Notably, this genogroup exhibited increased expression of VM proteins, a family of secreted AB-like toxins restricted to P1+ *Leptospira* strains and among the most highly induced genes during *in vivo* infection^36,46,56^. As VM proteins promote tissue invasion and systemic dissemination through the disruption of human cell-cell junctions, their selective upregulation in G4-Ict suggests that differential activity of these toxins contribute to pathogenicity, influencing bacterial dissemination and variation in virulence among genogroups. By contrast, G13-Pom exhibited reduced expression of virulence genes but induce the SOS response, consistent with a survival-oriented strategy under harmful conditions rather than one focused on infection. These results challenge the traditional view that host adaptation is primarily driven by gene acquisition and emphasize the importance of integrating multi-omics data to uncover the functional basis of virulence evolution in zoonotic pathogens.

Infection using mammalian macrophages demonstrates that *L. interrogans* genogroups diverge in their interactions with hosts. We identified distinct genogroup- and host-dependent immune response signatures, where G13-Pom elicited stronger pro-inflammatory responses in both human and murine macrophages than G4-Ict. One possible interpretation is that the two genotypes differ in their level of visibility to the innate immune surveillance. The broader induction of inflammatory response by G13-Pom, particularly in human macrophages, suggests that G13-Pom expose a ligand repertoire, including the genogroup-specific O-antigen structures encoded by the *rfb* locus (**Supplementary Fig. 19**) and other membrane-exposed proteins, that is efficiently detected by host cells. Conversely, G4-Ict has evolved mechanisms to mask these immunostimulatory determinants, an important feature for host dissemination^36,57,58^. This is particularly relevant given that leptospiral LPS is differentially recognized by Toll Like Receptor 2 (TLR2) and TLR4 murine and human cells^57,59^. Together, these findings support the idea that successful reservoir-associated leptospires limit excessive inflammation, and that the attenuated inflammatory profile of G4-Ict reflects an adaptive balance between transmission, persistence, and host damage. In addition, consistent with the higher prevalence in human infection, G4-Ict exhibited a markedly higher capacity to disrupt human epithelial cell-cell junction than G13-Pom, indicating an enhanced ability to breach physical barriers and facilitate tissue colonization. This mechanistic advantage for G4-Ict correlates with the higher expression of VM proteins which are involved in this disruption. Our findings underscore that genogroup-specific differences in immune recognition and epithelial barrier disruption shape *L. interrogans* pathogenic diversities, linking immune evasion with enhanced tissue dissemination for G4-Ict.

Collectively, these results demonstrate that host adaptation has profoundly structured the evolutionary diversification of the cosmopolitan species *L. interrogans*, generating genogroup-specific differences in genome plasticity, gene regulation, immune evasion and tissue dissemination. Beyond refining our understanding of leptospiral biology, this work highlights the importance of considering intraspecies diversity when investigating the mechanisms underlying virulence, host tropism and the emergence of zoonotic pathogens.

## Methods

### Ethics statement

Collection of the strains was conducted according to the Declaration of Helsinki. A written informed consent from patients was not required as the study was conducted as part of routine surveillance of the French National Reference Center for leptospirosis (Institut Pasteur, Paris, France) and no additional clinical specimens were collected for the purpose of the study. Cultures originating from human samples were anonymized.

### Bacterial strains and culture conditions

*Leptospira interrogans* strains used in this study for *in vitro* experiments are indicated in **Supplementary Table 5**. *Leptospira* strains were cultivated aerobically in Ellinghausen-McCullough-Johnson-Harris liquid medium (EMJH) at 30°C with shaking at 100 rpm or onto 1% agar solid EMJH media at 30°C for one month. To mimic host environmental condition, *Leptospira* strains were cultivated in EMEM medium (Sigma) supplemented with 5% rabbit serum and 0.72 nM FeSO4 at 37°C under 5% CO2, as previously described^43^. For all experiments, exponentially growing cultures of *Leptospira* were used.

### Genome dataset

Genome of the 368 *L. interrogans* isolates were obtained from the collection of the National Reference Centre for Leptospirosis (Institut Pasteur, Paris, France), which is a globally representative strain collection of isolates from animal and human samples gathered in the last 56 years (https://bigsdb.pasteur.fr/leptospira/).

Genomic DNA from *L. interrogans* strains used experimentally in this study (Supplementary Table 5) was extracted (Qiagen) and the whole-genome sequencing was performed using Oxford Nanopore Technology. Sequencing libraries were prepared following the SQK-RBK114-96 kit protocol and loaded onto a FLO-PRO114M flow cell for sequencing on a PromethION 2 Solo instrument. Sequencing runs were monitored using MinKNOW v24.06.10 and continued until a minimum genome coverage of 100x was achieved for each isolate. Basecalling was performed concurrently with sequencing using Dorado v0.7.4 in super-accurate mode. Reads were subsequently demultiplexed, adapter-trimmed and quality-filtered using a minimum quality score threshold of 10. De novo genome assemblies were generated with Flye v2.9.3^60^. Assembly quality was evaluated using Quast v5.2.0^61^ to check contigs size, BUSCO v5.4^62^ for genome completeness, Dfast v1.2.15^63^ for pseudogenes prediction and Bandage v0.9.0^64^ to confirmed of chromosome circularization.

### CgSNPs phylogenetic tree

A total of 368 genome sequences were subjected to comparative genomic analysis. All genomes were aligned against the UP-MMC-NIID LP strain reference genome (GenBank accession number GCA_001047635.1), and the core genome single nucleotide polymorphisms (cgSNPs) was performed using Snippy version 4.6.0. Recombinant regions were detected (included the entire *rfb* locus manually) and filtered from the cgSNPs alignments using Gubbins version 3.2.0^65^. A maximum-likelihood tree from the alignments was obtained using IQ-TREE version 2.4.0^66^ under the best-fit model of evolution (GTR+F+G4). Branch supports were assessed with bootstrap of 20,000 replicates. The outgroup used for tree construction was *L. kirschneri* 200702274 strain (the closer *Leptospira* species from *L. interrogans* in the phylogenetic species tree; GenBank accession number GCA_000244515.3). The tree was visualized using FigTree version 1.4.4. The genogroups were obtained from the cgSNPs phylogenetic tree and defined on the length phylogenetic distance method using TreeCluster version 1.0.3^67^.

### Pangenome openness and nucleotide diversity

Rarefaction curves of pan genome and core genome size estimations of each genogroup were obtained using a R script previously published^68^. Through the matrix of orthologous groups generated, the number of core genes and the cumulative number of different genes was calculated as the number of genomes sampled increased to obtain the mean core and pan genome size. Randomized calculations were used with 100 repetitions regarding the order of the genomes analyzed to obtain standard errors. Pangenome openness (γ) was calculated to assess the rate at which new genes are discovered with the addition of genomes. The openness parameter was estimated using the formula: γ =log(P/C) / log(N), where P is the total pangenome size, C is the core genome size, and N is the number of genomes analyzed. This approach quantifies the openness of the pangenome, with higher γ values indicating a more open pangenome architecture. Calculations were performed using RStudio v.2022.12.0. To measure genetic variation within a genogroup, nucleotide diversity was calculated for all coding sequences using the python library DendroPy v.4. For each gene, aligned nucleotide sequences were used to compute the average number of nucleotide differences per site between all pairs of sequences.

### Pfam domains analysis

Protein sequences from each genome were scanned for conserved domains using the Pfam-A HMM database and HMMER version 3.3.2 with an E-value threshold of 1e-5. Pfam counts were filtered to select only high-variance features (top 25% by variance across genomes). Remaining Pfam domains were normalized to relative abundance per genome.

To identify domains overrepresented in specific genogroups, a presence/absence binary matrix was constructed. For each genogroup, a one-sided Fisher’s exact test was applied to compare domain prevalence in the interested genogroup versus all other genomes. *p*-values were adjusted for multiple comparisons using the Benjamini-Hochberg method.

### SNP-enriched genes in *L. interrogans* isolates

All SNPs detected in *L. interrogans* isolates were filtered to retain only SNPs present inside open reading frame. Number of SNPs per gene was determined and the nomenclature of *L. interrogans* serovar Copenhageni strain Fiocruz L1-130 was used to perform COG and KEGG enrichment analysis.

### Positive selection and functional enrichment analysis for *L. interrogans* genogroups

*L. interrogans* genomes used in this study were annotated using Prokka (v.1.14.5). For each genogroups, orthologous groups were inferred based on protein sequences using OMA (v.2.5.0)^69^. Orthologous groups without paralogues which belong to the core genes were selected and transcript sequences were replaced to the protein sequences. To investigate signatures of positive selection across orthologous gene groups, we applied codon-based models implemented in the codeml program of the PAML package (v.4.9)^70^. Protein-coding sequences were aligned using Clustal Omega (v.1.2.4), and corresponding codon alignments were generated with PAL2NAL (v.14.1) using the "-nogap" option to maintain codon integrity. Phylogenetic trees, previously inferred from orthogroup gene trees, were used to constrain codeml analyses under user-specified topologies (runmode = 0). We focused on comparing site models M7 (beta distribution of ω across sites, disallowing ω > 1) and M8 (beta distribution with an additional class allowing ω > 1) to detect positive selection acting on a subset of sites. Likelihood ratio tests (LRTs) were conducted by extracting log-likelihood values from codeml output files, calculating test statistics as twice the difference in log-likelihoods between models, and assessing significance using the chi-square distribution with appropriate degrees of freedom. Orthogroups with significant M7 vs. M8 comparisons were further examined for sites under selection using the Naïve Empirical Bayes (NEB) approach. To ensure robustness, we also performed complementary model comparisons (M0 vs. M1a and M1a vs. M2a) and identified orthogroups with elevated ω values indicative of selection under the M0 model.

### Protein complex prediction and interface energy analysis

Protein-protein complex structures (LigB12-Elastin HTE20; LenA-Laminin β N-domain) were modeled using Alphafold 3 Multimer (v.2025.07.05; https://alphafoldserver.com/). The HTE-20 domain from elastin of human (UniProt: P15502), rat (UniProt: Q99372), bovine (UniProt: P04985) and pig (UniProt: A0A097ZMY9) were determined from sequence alignment using Clustal-Omega (v.1.2.4). The Laminin β N-domain from human (UniProt: P07942), rat (UniProt: D3ZQN7), bovine (UniProt: F1MNT4) and pig (UniProt: A0A5G2QKZ6) were determined from sequence alignment such as describe for elastin. The top-ranked model was selected for downstream analyses. The resulting mmCIF files were converted to PDB format using Gemmi (v.0.7.4). To determine the energy of interaction (ΔG interaction), structural models were processed using FoldX AnalyseComplex (v.2026.12.31) and Rosetta InterfaceAnalyzer suite (v.3.14). Negative ΔG interaction values indicate energetically favorable interfaces. Structural visualizations were performed using Mol* Viewer (v.5.6.0).

### RNA extraction, library preparation and sequencing of *L. interrogans*

Exponential phase cultures of five and four G4-Ict and G13-Pom strains, respectively, in EMJH and EMEM medium at 3 days post-incubation (at least in duplicate, **Supplementary Table 5**) were resuspended in QIAzol lysis reagent (Qiagen). Total RNAs were extracted using RNeasy Mini Kit (Qiagen), with a on column DNAse digestion step (Qiagen). The quality of all RNA samples was evaluated using an Agilent 2100 bioanalyzer (Agilent Technologies) to ascertain a RNA integrity number (RIN) higher than 9.

Sequencing Libraries were constructed using an Illumina Stranded Total RNA Prep Ligation using the Ribo-Zero Plus kit (Illumina, USA) with custom primer design on rRNA following the supplier’s recommendations. RNA sequencing was performed with the Illumina NextSeq 2000 for a target of 10M reads/sample. The RNA-seq analysis was performed with Sequana^71^ using the RNA-seq pipeline 0.19.2 (https://github.com/sequana/sequana_rnaseq) built on top of Snakemake 7.25.0^72^. Briefly, reads were trimmed from adapters using Fastp 0.23.2 then mapped to the *L. interrogans* G4 or G13 strains using Bowtie2^73^ and STAR^74^. FeatureCounts 2.0.1^75^ was used to produce the count matrix, assigning reads to features using the corresponding annotation from *L. interrogans* genomes using Prokka version 1.14.5. To identify differentially regulated genes between G4 and G13 strains, the core genes between the two genogroups were inferred based on protein sequences using OMA version 2.5.0^69^. In total, 3,052 core genes were found in all strains used, with 131 and 215 specific genes for genogroup G13 and G4, respectively. Statistical analysis on the count matrix was performed to identify differentially regulated genes. Clustering of transcriptomic profiles were assessed using a Principal Component Analysis (PCA). Differential expression testing was conducted using DESeq2 library 1.34.0^76^ scripts indicating the significance (Benjamini-Hochberg adjusted *p*-values, false discovery rate *p*.adj <0.05) and the effect size (fold-change) for each comparison.

### Proteomic analyses

#### Sample preparation

Bacterial proteins were extracted by repeated freeze–thaw lysis in ammonium bicarbonate (Lot: BCBR0615; Sigma-Aldrich) and sodium deoxycholate (Lot: SLBT7409; Sigma-Aldrich) buffer, followed by incubation on ice and centrifugation to remove cellular debris. Approximately 50 µg of protein from each sample was reduced with 10 mM dithiothreitol (DTT) to break cysteine disulfide bonds, alkylated with 40 mM iodoacetamide (IAA) to prevent disulfide bond reformation, and digested overnight with trypsin (Promega, Madison, USA). Formic acid was then added to quench digestion and precipitate detergent prior to LC–MS/MS analysis. Tryptic peptides were purified by centrifugation and solid-phase extraction using Oasis HLB plates (Waters, Milford), eluted with acetonitrile/formic acid, frozen, and lyophilized. Peptide concentrations were then confirmed using a colorimetric peptide assay prior to LC-MS/MS preparation. Finally, approximately 1 µg of peptides was loaded onto Evotip Pure C18 trap columns, which were preconditioned, equilibrated, washed, and maintained wet for subsequent mass spectrometry analysis.

#### Liquid chromatography-tandem mass spectrometry analysis

Peptides were analyzed on an Evosep One liquid chromatography (Evosep Biosystems) coupled to an Orbitrap FusionTM TribridTM (ThermoScientific) mass spectrometer equipped with a Nanospray FlexTM NG ion source (ThermoScientific). Peptides were separated using an endurance (EV1106) analytical column (ReproSil-Pur C18, 1.9 µm beads by Dr Maisch, 15 cm x 150 µm) with the extended method (15 samples per day) according to the manufacturer’s protocol. Briefly, in a 93-minute cycle time, peptides were injected into the mass spectrometer using a stainless-steel emitter (EV1086), ID 30 µm. The analytical column was equilibrated at 1500 nL/min, gradient flow at 220 nL/min for 88 min and then ramped up to 1500 nL/min for washing. Data were acquired in positive ion mode using a data-dependent acquisition (DDA) strategy with a 3-second cycle time. Full MS scans were collected in the Orbitrap at a resolution of 120,000 over a mass range of m/z 350–1800. The AGC target was set to 4.0e5 with a maximum injection time of 50 ms. The top precursors were selected for fragmentation using higher-energy collisional dissociation (HCD) and analyzed at a rapid scan rate in the ion trap. Fragmentation was performed with normalized collision energies of 28%, 30%, and 32%, using a quadrupole isolation window of 1.6 m/z. Only precursor ions with charge states from 2+ to 6+ were selected. Dynamic exclusion was enabled with a repeat count of 2, an exclusion duration of 7.5 s, and a mass tolerance of ±10 ppm. Lock mass correction was applied using a polysiloxane ion at m/z 445.12002.

#### Protein identification and quantification

Raw mass spectrometry data were processed using FragPipe (v24.0), incorporating MSFragger (v4.2)^77^ for database searching and IonQuant (v1.11.9) for label-free quantification. The search was performed against *a Leptospira interrogans* proteome database (uniprotkb_taxonomy_id_189518, downloaded on 2025-04-07, containing 3679 entries), and supplemented with common contaminant and decoy sequences. Mass tolerances were set to ±10 ppm for precursor ions and 20 ppm for fragment ions. Enzyme specificity was set to trypsin, allowing up to two missed cleavages. Carbamidomethylation of cysteine was used as a fixed modification, while oxidation of methionine and N-terminal acetylation were set as variable modifications. Peptide-spectrum matches were validated using Percolator (v3.7.1)^78^, and protein-level false discovery rate control at 1% using ProteinProphet. IonQuant performed intensity-based label-free quantification using the MaxLFQ algorithm with match-between-runs enabled.

### Macrophages and epithelial cells infection

THP-1 cells (a human monocyte cell line; ATCC TIB-202) and Raw 264.7 cells (a murine macrophage cell line) were cultured in RPMI 1640 medium (Gibco) supplemented with 10% heat-inactivated fetal bovine serum (Sigma) and 2 mM L-glutamine (Gibco). THP-1 cells were differentiated/activated into macrophages by a treatment with 50 nM PMA for 2 days following by a 24 hr incubation without PMA. Macrophages were infected with *L. interrogans* at a multiplicity of infection (MOI) of 50 bacteria-per-cell during 6h. Cells were incubated in 5% CO_2_ at 37°C.

Human epithelial cells (HEK-293T cells) were cultured in Minimum Essential Medium Eagle (Sigma) supplemented with 10% heat-inactivated fetal bovine serum (Sigma) and 2 mM L-glutamine (Gibco). Prior infection, *L. interrogans* strains were cultivated in EMEM medium (Sigma) supplemented with 5% rabbit serum and 0.72 nM FeSO_4_ at 37°C under 5% CO_2_ for 3 days and then diluted in the complete cell culture media prior the infection, at a multiplicity of infection of 50:1 during 24 hr. Cells were incubated in 5% CO_2_ at 37°C.

### Indirect immunofluorescence of epithelial cells

Epithelial cells were cultured on glass coverslips (SPL) coated with L-lysine (Sigma) at a concentration of 0.01% in water for a period of 40 min at 37°C. The coverslips were rinsed twice with PBS to remove excess of L-lysine, and the cells were seeded. At the indicated time points after infection, cell membranes were stained with FM4-64FX (5 µg/mL; ThermoFisher) and then cells were fixed with 4% paraformaldehyde for 15 min at room temperature (RT) and subsequently incubated for 10 min in 0.5% saponin (Sigma) in PBS and for 1 hr in 1% BSA (Sigma) and 0.075% saponin in PBS. The cells were incubated overnight at 4°C with the anti-pan-cadherin (PA5-16766; ThermoFisher) antibodies at 1:100. Cells were washed and incubated for 1 hr with Alexa Fluor 488 antibody (ThermoFisher) at 1:500. The nuclei were then stained with DAPI (1 µg/mL; ThermoFisher) for 10 min and mounted on a glass side using Fluoromount mounting medium (ThermoFisher). Fluorescence was analyzed using a Leica TCS SP8 Confocal System and the quantification was performed using Icy software (Version 3.0). For each experimental condition, at least 3 random fields and 100 cells were analyzed.

#### RNA extraction, library preparation and sequencing of human and murine macrophages

Total RNAs from human and murine macrophages at 6 hr PI (infected or not by five different *L. interrogans* G4-Ict and four different *L. interrogans* G13-Pom strains as described above) were extracted using QIAzol lysis reagent (Qiagen) and purified using RNeasy mini kit (Qiagen) with a DNAse treatment. Three replicates were used for uninfected conditions and two replicates for each *L. interrogans* strain used. cDNA libraries were prepared and were sequenced on an NovaSeq X Plus using Plasmidsaurus RNA-Seq service. STAR aligner v2.7.11 was used to map RNA-seq reads to the hg38 reference genome or GRCm39 reference genome for human and murine samples, respectively. Differential expression analysis was performed using edgeR v4.0.16 with filtering for low-expressed genes with edgeR::filterByExpr (default values). Functional enrichment for human and mouse samples was performed using GSEApy v0.12 using the MSigDB Hallmark gene set.

## DATA AVAILABILITY

Sequenced genomes generated in this study have been deposited in the NCBI database under BioProject accession codes PRJNA1471134. The raw fastq files of RNA-sequencing have been deposited in NCBI’s Gene Expression Omnibus and are accessible through GEO Series accession number GSE311287 and GSE330072, for *L. interrogans* and mammal hosts, respectively. The mass spectrometry proteomics data have been deposited to the ProteomeXchange Consortium via the PRIDE partner repository with the dataset identifier PXD078695.

## ACKNOWLEDGEMENTS

This work has received financial support by the National Institutes of Health grant P01 AI 168148 (MP). We thank Jean-François Mariet of the French National Reference Center for Leptospirosis for curation of BIGSdb. We also acknowledge L. Lemée and G. Haustant from Biomics Platform, C2RT, Institut Pasteur, Paris, France, supported by France Génomique (ANR-10-INBS-09) and IBISA. HP acknowledges support from the National Science and Engineering Research Council of Canada (NSERC) Discovery Grant RGPIN-2026-04712 and the Early Career Supplement DGECR-2026-00389. The work at the University of Victoria Genome BC Proteomics Centre was supported by the BC Proteomics Center (BCPC) from Genome British Columbia for operations and technology development (374PRO and 384 PRO). The Authors would like to thank Angela M. Jackson for assistance with the sample preparation for proteomics analysis. We are grateful to K. Coullin and M. Lago for critically reading the manuscript. The funders had no role in study design, data collection and analysis, decision to publish, or preparation of the manuscript.

## Notes

### Competing Interest Statement

The authors have declared no competing interest.

## REFERENCES

1. Woolhouse, M. E. J. & Gowtage-Sequeria, S. Host Range and Emerging and Reemerging Pathogens. Emerg. Infect. Dis. 11, 1842–1847 (2005).

2. Jones, K. E. et al. Global trends in emerging infectious diseases. Nature 451, 990–993 (2008).

3. Plowright, R. K. et al. Pathways to zoonotic spillover. Nat. Rev. Microbiol. 15, 502–510 (2017).

4. Costa, F., et al. Global Morbidity and Mortality of Leptospirosis: A Systematic Review. *PLoS Negl. Trop. Dis*. 9, e0003898 (2015).

5. Mwachui, M. A., Crump, L., Hartskeerl, R., Zinsstag, J. & Hattendorf, J. Environmental and Behavioural Determinants of Leptospirosis Transmission: A Systematic Review. PLoS Negl. Trop. Dis. 9, e0003843 (2015).

6. Halliday, J. E. B. et al. Zoonotic causes of febrile illness in malaria endemic countries: a systematic review. Lancet Infect. Dis. 20, e27–e37 (2020).

7. Pijnacker, R. et al. Marked increase in leptospirosis infections in humans and dogs in the Netherlands, 2014. Eurosurveillance 21, 30211 (2016).

8. Beauté, J. et al. Epidemiology of reported cases of leptospirosis in the EU/EEA, 2010 to 2021. Eurosurveillance 29, 2300266 (2024).

9. Ellis, W. A. Animal Leptospirosis. in Leptospira and Leptospirosis (ed. Adler, B.) 99–137 (Springer, Berlin, Heidelberg, 2015). doi:10.1007/978-3-662-45059-8_6.

10. Hagedoorn, N. N. et al. Global distribution of Leptospira serovar isolations and detections from animal host species: a systematic review and online database. Trop. Med. Int. Health TM IH 29, 161–172 (2024).

11. Rajapakse, S., Fernando, N., Dreyfus, A., Smith, C. & Rodrigo, C. Leptospirosis. Nat. Rev. Dis. Primer 11, 1–19 (2025).

12. Picardeau, M. Virulence of the zoonotic agent of leptospirosis: still terra incognita? Nat. Rev. Microbiol. 15, 297–307 (2017).

13. Giraud-Gatineau, A. et al. Shaping the future of Leptospira serotyping. J. Med. Microbiol. 74, 002059 (2025).

14. Davignon, G. et al. Leptospirosis: toward a better understanding of the environmental lifestyle of Leptospira. *Front*. Water 5, (2023).

15. Ghosh, S. et al. Transcriptional diversification in a human-adapting zoonotic pathogen drives niche-specific evolution. Nat. Commun. 16, 2067 (2025).

16. Hafner, L. et al. Differential stress responsiveness determines intraspecies virulence heterogeneity and host adaptation in Listeria monocytogenes. Nat. Microbiol. 9, 3345–3361 (2024).

17. Vincent, A. T., et al. Revisiting the taxonomy and evolution of pathogenicity of the genus Leptospira through the prism of genomics. PLoS Negl. Trop. Dis. 13, e0007270 (2019).

18. Chinchilla, D., et al. Phylogenomics of Leptospira santarosai, a prevalent pathogenic species in the Americas. PLoS Negl. Trop. Dis. 17, e0011733 (2023).

19. Christova, I., Tasseva, E. & Manev, H. Human leptospirosis in Bulgaria, 1989-2001: epidemiological, clinical, and serological features. Scand. J. Infect. Dis. 35, 869–872 (2003).

20. Tubiana, S. et al. Risk Factors and Predictors of Severe Leptospirosis in New Caledonia. PLoS Negl. Trop. Dis. 7, e1991 (2013).

21. Shintaku, M., Itoh, H. & Tsutsumi, Y. Weil’s disease (leptospirosis) manifesting as fulminant hepatic failure: report of an autopsy case. Pathol. Res. Pract. 210, 1134–1137 (2014).

22. Hu, W., Lin, X. & Yan, J. Leptospira and leptospirosis in China. Curr. Opin. Infect. Dis. 27, 432–436 (2014).

23. Hochedez, P. et al. Factors Associated with Severe Leptospirosis, Martinique, 2010-2013. Emerg. Infect. Dis. 21, 2221–2224 (2015).

24. Grillová, L., et al. Circulating genotypes of Leptospira in French Polynesia : An 9-year molecular epidemiology surveillance follow-up study. PLoS Negl. Trop. Dis. 14, e0008662 (2020).

25. Eves, C., Kjelsø, C., Benedetti, G., Jørgensen, C. S. & Krogfelt, K. A. Trends in human leptospirosis in Denmark, 2012-2021. Front. Cell. Infect. Microbiol. **13**, (2023).

26. Mišić-Majerus, L., Kišek, T. C. & Ružić-Sabljić, E. Leptospirosis and characterization of Leptospira isolates from patients in Koprivnica-Križevci County, Croatia from 2000–2004. Access Microbiol. 5, acmi000431 (2023).

27. Ferreira, L. C., Filho, L. de F. F., Cosate, M. R. V. & Sakamoto, T. Genetic structure and diversity of the rfb locus of pathogenic species of the genus Leptospira. Life Sci. Alliance 7, (2024).

28. Liao, J. et al. Nationwide genomic atlas of soil-dwelling Listeria reveals effects of selection and population ecology on pangenome evolution. Nat. Microbiol. 6, 1021–1030 (2021).

29. Amamura, T. A. et al. Proteolytic activity of secreted proteases from pathogenic leptospires and effects on phagocytosis by murine macrophages. Microbes Infect. 27, 105469 (2025).

30. Eshghi, A. et al. An extracellular Leptospira interrogans leucine-rich repeat protein binds human E- and VE-cadherins. Cell. Microbiol. 21, e12949 (2019).

31. Otto, H. et al. The spvB gene-product of the Salmonella enterica virulence plasmid is a mono(ADP-ribosyl)transferase. Mol. Microbiol. 37, 1106–1115 (2000).

32. Colclough, A. L., Scadden, J. & Blair, J. M. A. TetR-family transcription factors in Gram-negative bacteria: conservation, variation and implications for efflux-mediated antimicrobial resistance. BMC Genomics 20, 731 (2019).

33. Tomar, D. et al. MCUR1 Is a Scaffold Factor for the MCU Complex Function and Promotes Mitochondrial Bioenergetics. Cell Rep. 15, 1673–1685 (2016).

34. Wang, S. et al. Revealing roles of S-layer protein (SlpA) in Clostridioides difficile pathogenicity by generating the first slpA gene deletion mutant. Microbiol. Spectr. 12, e0400523 (2024).

35. Gupta, A. et al. MarR Family Transcription Factors from Burkholderia Species: Hidden Clues to Control of Virulence-Associated Genes. Microbiol. Mol. Biol. Rev. MMBR 83, e00039–18 (2018).

36. Giraud-Gatineau, A. et al. Evolutionary insights into the emergence of virulent Leptospira spirochetes. PLOS Pathog. 20, e1012161 (2024).

37. Silva, L. P. et al. Evaluation of two novel leptospiral proteins for their interaction with human host components. Pathog. Dis. 74, ftw040 (2016).

38. Atzingen, M. V. et al. Lp95, a novel leptospiral protein that binds extracellular matrix components and activates e-selectin on endothelial cells. J. Infect. 59, 264–276 (2009).

39. Abramson, J. et al. Accurate structure prediction of biomolecular interactions with AlphaFold 3. Nature 630, 493–500 (2024).

40. Hinds, A. et al. AlphaFold reveals how pathogenic Leptospira use cross-kingdom thiol-disulfide exchange to evade the complement membrane attack complex. mBio 0, e00878–26 (2026).

41. Lin, Y.-P. et al. Repeated Domains of Leptospira Immunoglobulin-like Proteins Interact with Elastin and Tropoelastin. J. Biol. Chem. 284, 19380–19391 (2009).

42. Barbosa, A. S. et al. A Newly Identified Leptospiral Adhesin Mediates Attachment to Laminin. Infect. Immun. 74, 6356–6364 (2006).

43. Garcia, L. E., et al. DMEM and EMEM as alternate growth media for pathogenic Leptospira. PLoS Negl. Trop. Dis. 20, e0014136 (2026).

44. Haake, D. A. & Matsunaga, J. Leptospiral Immunoglobulin-Like Domain Proteins: Roles in Virulence and Immunity. Front. Immunol. 11, (2021).

45. Zhu, W. et al. MPL36, a major plasminogen (PLG) receptor in pathogenic Leptospira, has an essential role during infection. PLOS Pathog. 19, e1011313 (2023).

46. Giraud-Gatineau, A., Haustant, G., Monot, M., Picardeau, M. & Benaroudj, N. In vivo dual RNA-Seq uncovers key effectors of epithelial barrier disruption by an extracellular pathogen. Nat. Commun. 17, 2274 (2026).

47. Weinert, L. A. et al. Genomic signatures of human and animal disease in the zoonotic pathogen Streptococcus suis. Nat. Commun. 6, 6740 (2015).

48. Strydom, H. et al. Evaluating sub-typing methods for pathogenic Yersinia enterocolitica to support outbreak investigations in New Zealand. Epidemiol. Infect. 147, e186 (2019).

49. Rosconi, F. et al. A bacterial pan-genome makes gene essentiality strain-dependent and evolvable. Nat. Microbiol. 7, 1580–1592 (2022).

50. Santos, L. A. et al. Genomic Comparison Among Global Isolates of L. interrogans Serovars Copenhageni and Icterohaemorrhagiae Identified Natural Genetic Variation Caused by an Indel. Front. Cell. Infect. Microbiol. 8, (2018).

51. Dekker, J. P. Within-Host Evolution of Bacterial Pathogens in Acute and Chronic Infection. Annu. Rev. Pathol. Mech. Dis. 19, 203–226 (2024).

52. Didelot, X., Walker, A. S., Peto, T. E., Crook, D. W. & Wilson, D. J. Within-host evolution of bacterial pathogens. Nat. Rev. Microbiol. 14, 150–162 (2016).

53. Giraud-Gatineau, A. et al. Inter-species Transcriptomic Analysis Reveals a Constitutive Adaptation Against Oxidative Stress for the Highly Virulent Leptospira Species. Mol. Biol. Evol. 41, msae066 (2024).

54. Brown, S. P., Cornforth, D. M. & Mideo, N. Evolution of virulence in opportunistic pathogens: generalism, plasticity, and control. Trends Microbiol. 20, 336–342 (2012).

55. Sokurenko, E. V., Gomulkiewicz, R. & Dykhuizen, D. E. Source–sink dynamics of virulence evolution. Nat. Rev. Microbiol. 4, 548–555 (2006).

56. Chaurasia, R., Marroquin, A. S., Vinetz, J. M. & Matthias, M. A. Pathogenic Leptospira Evolved a Unique Gene Family Comprised of Ricin B-Like Lectin Domain-Containing Cytotoxins. Front. Microbiol. 13, (2022).

57. Werts, C. et al. Leptospiral lipopolysaccharide activates cells through a TLR2-dependent mechanism. Nat. Immunol. 2, 346–352 (2001).

58. Bonhomme, D. & Werts, C. Host and Species-Specificities of Pattern Recognition Receptors Upon Infection With Leptospira interrogans. Front. Cell. Infect. Microbiol. 12, 932137 (2022).

59. Nahori, M.-A. et al. Differential TLR Recognition of Leptospiral Lipid A and Lipopolysaccharide in Murine and Human Cells1. J. Immunol. 175, 6022–6031 (2005).

60. Kolmogorov, M., Yuan, J., Lin, Y. & Pevzner, P. A. Assembly of long, error-prone reads using repeat graphs. Nat. Biotechnol. 37, 540–546 (2019).

61. Gurevich, A., Saveliev, V., Vyahhi, N. & Tesler, G. QUAST: quality assessment tool for genome assemblies. Bioinformatics 29, 1072–1075 (2013).

62. Simão, F. A., Waterhouse, R. M., Ioannidis, P., Kriventseva, E. V. & Zdobnov, E. M. BUSCO: assessing genome assembly and annotation completeness with single-copy orthologs. Bioinformatics 31, 3210–3212 (2015).

63. Tanizawa, Y., Fujisawa, T. & Nakamura, Y. DFAST: a flexible prokaryotic genome annotation pipeline for faster genome publication. Bioinformatics 34, 1037–1039 (2018).

64. Wick, R. R., Schultz, M. B., Zobel, J. & Holt, K. E. Bandage: interactive visualization of de novo genome assemblies. Bioinformatics 31, 3350–3352 (2015).

65. Croucher, N. J. et al. Rapid phylogenetic analysis of large samples of recombinant bacterial whole genome sequences using Gubbins. Nucleic Acids Res. 43, e15 (2015).

66. Nguyen, L.-T., Schmidt, H. A., von Haeseler, A. & Minh, B. Q. IQ-TREE: A Fast and Effective Stochastic Algorithm for Estimating Maximum-Likelihood Phylogenies. Mol. Biol. Evol. 32, 268–274 (2015).

67. Balaban, M., Moshiri, N., Mai, U., Jia, X. & Mirarab, S. TreeCluster: Clustering biological sequences using phylogenetic trees. PLOS ONE 14, e0221068 (2019).

68. Méric, G. et al. A Reference Pan-Genome Approach to Comparative Bacterial Genomics: Identification of Novel Epidemiological Markers in Pathogenic Campylobacter. PLOS ONE 9, e92798 (2014).

69. Dylus, D. et al. How to build phylogenetic species trees with OMA. F1000Research 9, 511 (2022).

70. Yang, Z. PAML 4: Phylogenetic Analysis by Maximum Likelihood. Mol. Biol. Evol. 24, 1586–1591 (2007).

71. Cokelaer, T., Desvillechabrol, D., Legendre, R. & Cardon, M. ‘Sequana’: a Set of Snakemake NGS pipelines. J. Open Source Softw. 2, 352 (2017).

72. Köster, J. & Rahmann, S. Snakemake—a scalable bioinformatics workflow engine. Bioinformatics 28, 2520–2522 (2012).

73. Langmead, B. & Salzberg, S. L. Fast gapped-read alignment with Bowtie 2. Nat. Methods 9, 357–359 (2012).

74. Dobin, A. et al. STAR: ultrafast universal RNA-seq aligner. Bioinformatics 29, 15–21 (2013).

75. Liao, Y., Smyth, G. K. & Shi, W. featureCounts: an efficient general purpose program for assigning sequence reads to genomic features. Bioinformatics 30, 923–930 (2014).

76. Love, M. I., Huber, W. & Anders, S. Moderated estimation of fold change and dispersion for RNA-seq data with DESeq2. Genome Biol. 15, 550 (2014).

77. Kong, A. T., Leprevost, F. V., Avtonomov, D. M., Mellacheruvu, D. & Nesvizhskii, A. I. MSFragger: ultrafast and comprehensive peptide identification in mass spectrometry–based proteomics. Nat. Methods 14, 513–520 (2017).

78. Käll, L., Canterbury, J. D., Weston, J., Noble, W. S. & MacCoss, M. J. Semi-supervised learning for peptide identification from shotgun proteomics datasets. Nat. Methods 4, 923–925 (2007).

