## Supplementary Information for "Host adaptation drives genome evolution and virulence diversification in a bacterial zoonotic pathogen"

### SUPPLEMENTARY METHODS

#### Determination of *rfb* locus gene orthogroups

A total of 368 *L. interrogans* genomes were annotated using Prokka version 1.14.5[1]. Orthogroups between all the protein sequence sets were inferred using the OrthoFinder version 2.5.5[2], applying a minimum threshold of 80% protein sequence identity to define orthologous relationships. To identity orthogroups associated with the *rfb* locus, a subset of genomes containing the *rfb* locus entirely located within a single contig was selected. The rfb locus was delineated by the genes encoding a MarR-family transcriptional regulator (*marR*) and a sodium/sulfate symporter (*sdcS*), as previously described[3]. Genes located between these two boundary markers were extracted, and their orthogroups were identified. These locus-defining orthogroups were then compared across fragmented genomes to include partially assembled *rfb* loci. This process resulted in the selection of 140 *rfb*-associated orthogroups across all *L. interrogans* genomes.

#### Hierarchical clustering of *L. interrogans* isolates on the *rfb* locus

The hierarchical clustering method was used to group isolates/orthogroups with similar profiles in the rfb locus. This process resulted in a binary matrix of 140 orthogroups by isolate, indicating presence (1) or absence (0) of each orthogroup per isolate. The binary matrix was processed in R version 4.2.2 where hierarchical clustering was performed using the Ward's minimum variance method. Dendrogram representing isolate clustering based on *rfb* locus were constructed using ape and stats packages. Heatmap was generated using the pheatmap package, displaying the distribution of *rfb* orthogroups across isolates. The resulting hierarchical structure allowed for grouping of isolates based on *rfb* gene content and was subsequently used to infer serogroup identity in isolates with unknown serological classification.

#### Human serum survival

Sera were obtained from healthy donors. The blood collection was carried out in accordance with the approved French Ministry of Research and French Ethics Committee protocols by the Etablissement Français du Sang (EFS, n°18/EFS/041). Written consent was received from all participants donating blood for the study. Exponentially growing *Leptospira* strains were centrifuged at 2,600 g for 15 min, washed three times and resuspended into PBS 1X (Gibco). *Leptospira interrogans* isolates were adjusted to $5x10^8$ leptospires/ml and incubated with 20% human serum or 20% heat-inactivated human serum diluted in PBS. For colony-forming unit (CFU) determination, Leptospira were diluted in PBS and plated on EMJH agar plates. Colonies were counted and the percent survival (% of CFU) was calculated as the ratio of CFU for bacteria incubated with human serum to bacteria incubated without human serum.

**SNP distribution across *L. interrogans* genome**

To analyze the distribution of SNPs across all *L. interrogans* genomes and assess their association with specific localization or gene length, we used the list of detected SNPs generated by snippy analysis and extract gene coordinates and the number of SNPs per gene. To evaluate whether SNP-enriched genes were randomly distributed across the genome, we performed a Kolmogorov-Smirnov (KS) test, comparing the positional distribution of SNP-containing genes to all annotated genes. Additionally, a circular genome representation was generated using Matplotlib, where SNP density was generated and plotted as bars around the genome circle to visually assess SNP clustering. Then, to determine if the observed SNP enrichment in specific genes was biased by gene length, we calculated Spearman's rank correlation between gene size and SNP count. Furthermore, we tested whether SNP-containing genes were significantly larger than non-SNP-containing genes using a Wilcoxon rank-sum test. All statistical analyses were conducted in Python using SciPy and StatsModels libraries.

**Gene expression by RT-qPCR**

Total RNAs from human and murine macrophages were extracted using QIAzol lysis reagent (Qiagen) and purified with RNeasy columns (Qiagen). Reverse transcription of mRNA to cDNA was carried out using the iScript cDNA Synthesis kit (Bio-rad), followed by cDNA amplification using the SsoFast EvaGreen Supermix (Bio-rad). All primers used in this study are listed in **Supplementary Table 12**. Reactions were performed using the CFX96 real-time PCR detection system (Bio-Rad). The relative gene expression was assessed according to the $2^{-\Delta Ct}$ method using *gadph* as reference gene for human and mouse macrophages. To perform comparative analysis between genogroups for each type of macrophages, expression values were normalized across samples for each gene across all samples (using z-score standardization; scikit-learn library version 1.4.2), facilitating comparative analysis across conditions. To analyze global patterns in gene expression by genogroup, principal component analysis (PCA) was performed on the normalized gene expression at 6h post-infection.

**Lipid A analysis by Matrix-Assisted Laser Desorption/Ionization (MALDI) mass spectrometry (MS)**

All solvents were MS grade and purchased from Fisher Chemical (Hampton, NH, USA). *Leptospira* lipid A was analyzed as previously described[4,5]. Bacterial pellets were resuspended in water (n=3 biological replicates/strain). One microliter of the resulting sample was dried on a stainless steel MALDI plate in technical triplicates (MFX µFocus plate 12×8 c 2,400 µm 0.7 T; Hudson Surface Technology, Inc., South Korea) and overlaid with one microliter of 0.2 M citric acid, 0.1 M sodium citrate buffer. The plate was then incubated for 30 min at 110°C in a humidity chamber, thoroughly rinsed with 50 ml of water, and dried. One microliter of norharmane matrix (modified to 7 mg/mL in 2:1 v/v chloroform and methanol) was spotted on top of each sample prior to MALDI MS analysis.

Data were collected in the negative ion mode over a m/z 600-2,500 scan range using a timsTOF flex MALDI-2 (Bruker, Bremen, Germany). The instrument was calibrated with Agilent Calibration mix (Agilent Technologies, Santa Clara, CA, USA) in an electrospray mode. The following settings were used for tandem MS analyses: m/z 200-2,500 scan range: shots/spot: 3,000, frequency: 5,000 Hz, collision RF: 4,000 Vpp, transfer time: 110 µs, prepulse storage: 11 µs, isolation width: m/z 2and collision energy: 130 eV. Data were analyzed using Compass Data Analysis v 6.1 (Bruker).

### SUPPLEMENTARY DATA

**Supplementary Fig. 1. Geographical distribution of *L. interrogans* genogroups**
World map illustrating the presence of each *L. interrogans* genogroup. Countries with isolates are shaded in black. Data by serogroup, hosts and geographic region were provided below each map when available.

**Supplementary Fig. 2. Hierarchical clustering of *rfb* locus in *L. interrogans* isolates**
Orthogroup inference was performed on the *rfb* locus across all *L. interrogans* isolates included in this study, using a protein identity cutoff of 80%. The heatmap displays presence/absence of *rfb*-associated orthogroups, with white indicating absence and grey indicating presence. Rows represent individual orthogroups, while columns correspond to distinct *L. interrogans* isolates. The dendrogram above the heatmap shows clustering of isolates based on orthogroup profiles. Below the heatmap, the serogroup identity of each isolate is indicated. Seven distinct clusters are highlighting, each correspond to specific serogroups, enabling serogroup prediction for untyped isolates.

**Supplementary Fig. 3. Genome diversity and pangenome dynamics in *L. interrogans***
(**a**) Distribution of total SNP counts within coding-sequences per *L. interrogans* genogroup. In the x-axis, the number of genomes belonged to the genogroup is indicated. Genogroups with the lowest SNP counts are highlighted in blue.
(**b**) Pangenome openness across genogroups. Genogroups with the lowest pangenome openness are highlighted in blue.
**(c)** Rarefaction and accumulation curves of core and pangenomes in *L. interrogans* genogroups. The mean core (blue) and the pangenome (in red) sizes are plotted for each genogroups by increasing number of genomes. Randomizing the order of genomes was repeated 100 times to obtain the average standard deviation. For each genogroup, the pangenome openness was indicated.

**Supplementary Fig. 4. Protein domain and associated-virulence genes diversity in *L. interrogans* genogroups**

Heatmap showing the distribution of selected Pfam domains and virulence genes across *L. interrogans* isolates, aligned with the cgSNPs tree.

**Supplementary Fig. 5. Pfam domains categories over- and under-represented in *L. interrogans* genogroups G4-Ict and G13-Pom**

Overrepresented (a) and underrepresented (b) Pfam domain categories in *L. interrogans* G4-Ict and G13-Pom. The number of Pfam domains present in each category was indicated in the x-axis.

**Supplementary Fig. 6. Genomic variation hotspots among *L. interrogans* isolates**

(**a**) Histogram showing the distribution of SNP counts across *L. interrogans* core-genes. The x-axis represents SNP count per gene, while the y-axis indicates the number of genes.

(**b**) Functional categorization of genes with ≥61 using COG classification (green: Cellular process and signaling, purple: Information storage and processing, orange: Metabolism). Sub-categories are detailed on the y-axis. Fisher's exact test with Benjamini-Hochberg method correction was used to estimate the enrichment of categories. *, $p<0.05$; **, $p<0.01$.

(**c**) Schematic representation of major functional categories enriched in genes with SNPs, annotated using the nomenclature of *L. interrogans* serovar Copenhageni Fiocruz L1-130 strain. The number of SNPs detected across *L. interrogans* isolates is indicated for each gene enriched in SNPs. Known virulence-associated genes are indicated in red.

(**d**) Proportion of genes under positive selection (dN/dS >1) within SNP-enriched categories across genogroups. Statistical significance determined by Fisher's exact test was used. *, $p<0.05$; **, $p<0.01$; ***, $p \leq 0.001$.

**Supplementary Fig. 7. Distribution of point mutation across *L. interrogans* genome**

(**a**) Distribution of point mutation on the chromosome 1 of *L. interrogans*. The circular plot displays the chromosome size of *L. interrogans* with ORFs in green and SNP density in black lines (window size: 1,000 bp). Kolmogorov-Smirnov (KS) test was used to evaluate whether SNP-enriched genes were randomly distributed across the genome (KS and *p*-value are indicated in the panel).

(**b**) Gene length distribution for SNP-enriched (>60 SNPs in blue) and non-enriched (≤ 60 SNPs in red) genes (left panel). Boxplot showing the distribution of gene lengths for SNP-enriched and non-enriched genes (right panel). Spearman's rank correlation was used to determine if the enrichment of SNPs in specific genes is associated with gene length and Wilcoxon rank-sum test was used to test whether genes containing SNP were significantly larger than genes that do not associate with SNPs (Spearman correlation and *p*-values were indicated in the panel).

**Supplementary Fig. 8. SNP distribution in virulence-associated genes in *L. interrogans***

Bar plot displaying SNP counts per virulence-associated gene, annotated using *L. interrogans* serovar Copenhageni strain Fiocruz L1-130 nomenclature.

**Supplementary Fig. 9. Functional categorization of conserved genes in *L. interrogans***

(**a**) COG classification of core genes with ≤1 SNP. Genes were divided into four groups (green: Cellular process and signaling, purple: Information storage and processing, orange: Metabolism, white: Poorly characterized).

(**b**) Sub-categories of COG annotations with corresponding gene counts. Fisher's exact test with Benjamini-Hochberg method correction was used to estimate the enrichment. *, $p$ <0.05; **, $p$ <0.01.

(**c**) KEGG pathway enrichment analysis of genes without substitution in *L. interrogans* isolates. The y-axis shows enriched pathways, and the x-axis shows the number of genes counts per pathway.

**Supplementary Fig. 10. Impact of SNP-enriched pathways on genogroup distribution and host specificity**

(**a**) Proportions and type of substitutions in SNP-enriched genes across genogroups. The x-axis represents the genogroup and the y-axis indicates substitutions counts. The nature of their substitutions is indicated in the legend.

(**b**) Neighbor-joining tree based on protein alignments of SNP-enriched genes. Colored triangles represent genogroup with serovar information annotated. The tree from the alignments was obtained using IQ-TREE under the best-fit model of evolution (LG+F+R8). Branch supports were assessed with bootstrap of 10,000 replicates.

**Supplementary Fig. 11. Confidence scores of predicted interactions between *Leptospira* LenA and Laminin proteins from different host species.**

Distribution of AlphaFold confidence scores (0.8 x ipTM + 0.2 x pTM) for predicted complexes between LenA from G4-Ict (blue) or from G13-Pom (yellow) and laminin domains from the α (a), β (b) and γ (c) subunits of human, rat, bovine and porcine hosts. Each point represents an independent AlphaFold model prediction. Data are the mean ± SD from 5 models.

**Supplementary Fig. 12. Growth kinetics of *L. interrogans* G4-Ict and G13-Pom**

Growth curves of *L. interrogans* G4-Ict and G13-Pom strains cultured in EMJH (bacterial medium) and in EMEM medium. Bacterial growth was enumerated using a Petroff-Hausser counting chamber. Strains identification and genogroup are indicated in the legend. Data are the mean ± SD from 3 independent experiments.

**Supplementary Fig. 13. Transcriptomic analysis of G4-Ict and G13-Pom in host-like condition compared to *in vitro* condition**

(**a**) Venn diagram showing differentially expressed genes (DEGs; *p*.adj ≤0.05; $|Log_2FC|$ >0) for genogroup G4-Ict (5 strains) and G13-Pom (4 strains) in EMEM medium at 37°C, 5% CO2 compared to EMJH medium at 30°C after 3 days of incubation. For upregulated (upper diagram) and downregulated (lower diagram) genes, the number of common and specific genes are indicated.

(**b**) The y-axis shows enriched pathways and values of the x-axis are the mean of $Log_2FC$ of DEGs for each enriched pathway. The blue and red circle correspond to the mean of $Log_2FC$ of G4-Ict and G13-Pom, respectively. Statistical significance determined by Fisher's exact test was used. *, *p* <0.05; **, *p* <0.01; ***, *p* ≤0.001.

(**c-d**) KEGG pathway enrichment analysis of DEGs specific to G4-Ict (c) and G13-Pom (d). The y-axis shows enriched pathways and values of the x-axis are the mean of $Log_2FC$ of DEGs for each enriched pathway. Statistical significance determined by Fisher's exact test was used. *, *p* ≤0.05; **, *p* ≤0.01; ***, *p* ≤0.001.

**Supplementary Fig. 14. Regulation and functional classification of differentially expressed genes specific to *L. interrogans* genogroups G4-Ict and G13-Pom**

(**a**) Percentage of upregulated and downregulated SNP-enriched genes related to Cell wall, membrane and Signal transduction mechanism categories in the genogroup G4-Ict and G13-Pom.

(**b**) Differential expression of genes exclusive to G4-Ict (blue) and G13-Pom (red) when cultured in EMEM medium compared to EMJH medium at 3 days post-incubation (*p*.adj ≤0.05). Both upregulated and downregulated genes are shown.

(**c-d**) Functional classification of G4-Ict or G13-Pom exclusive DEGs based on COG categories. Upregulated genes (c) and downregulated genes (d) in each genogroup are represented.

**Supplementary Fig. 15. Inter-genogroup transcriptional differences between *L. interrogans* G4-Ict and G13-Pom under host-like conditions.**

List of the 20 most up-regulated genes (*p*.adj ≤0.05) in genogroup G4-Ict compared to G13-Pom (left) and in genogroup G13-Pom compared to G4-Ict (right). Virulence-associated genes are indicated in red.

**Supplementary Figure 16. Genogroup-specific induction of innate immune responses in macrophages**

Normalized expression levels of inflammatory response genes in human (**a**) or murine (**b**) macrophages infected with *L. interrogans* strains from genogroups G4-Ict (n= 5 strains) and G13-Pom (n= 4 strains) at a multiplicity of infection (MOI) of 50:1 at 6 hr post-infection. Gene expression was analyzed using *gapdh* as a reference gene and expressed as $Log_2$(FC). Gradient color from blue to red indicates low to high $Log_2$(FC) values. Principal component analysis (PCA) was performed on normalized gene expression to assess genogroup-specific clustering. Variance explained by each principal component is indicated. Data represent the mean ± SD from three independent experiments.

**Supplementary Fig. 17. Resistance of *L. interrogans* genogroups G4-Ict and G13-Pom to human serum**

Each *L. interrogans* G4-Ict (n= 5 strains) and G13-Pom (n= 4 strains) strains were incubated in 20% of normal or heat-inactivated human serum for 2h. Living bacteria were enumerated by CFU, and the percentage survival was determined relative to CFU counts obtained in heat-inactivated human serum. *L. biflexa* strain was used as control for serum activity. Data are the mean ± SD from 3 independent experiments for each *Leptospira* strains used. Unpaired two-tailed Student's *t* test was used. ns: no significant.

**Supplementary Fig. 18. Lipid A analysis of *L. interrogans* genogroups G4-Ict and G13-Pom**

(**a**) Lipid A profiles, as determined by MALDI MS.

(**b**) Tandem MS analysis of the base lipid A ion, m/z 1748.26. Data are representative of three biological replicates and were acquired on a timsTOF flex MALDI-2 in the negative ion mode. r. int.: relative intensity.

**Supplementary Fig. 19. Genetic organization of the *rfb* locus in G4-Ict and G13-Pom strains**

The alignment of the *rfb* locus located between *marR* and *dass* genes was performed using the five G4-Ict strains and the four G13-Pom strains used in this study. Common, specific serogroup Icterohaemorrhagiae, specific serogroup Pomona genes are indicated in black, blue and red, respectively. Inter-variable genes, in orange, are indicated and present only in G13-Pom strains.

**Supplementary Table 1. Information of *L. interrogans* strains used in this study**

Details of 368 *L. interrogans* isolates are indicated. Information regarding genogroup, serogroup, serogroup prediction through *rfb* locus clustering, serovar, country, host isolated, and genome size are indicated.

**Supplementary Table 2. Pfam domains under-represented in *L. interrogans* genogroups**

List of Pfam domains significantly enriched in each genogroup based on presence/absence profiles. For each domain, a one-sided Fisher's exact test was performed comparing its frequency within the focal genogroup versus all other genogroups. *p*-values were corrected for multiple testing using the Benjamini–Hochberg method. Columns include domain-level gene counts, odds ratios, unadjusted and adjusted *p*-values, and percentage presence inside and outside of the genogroup.

**Supplementary Table 3. KEGG pathway enrichment analysis of genes under positive selection in each *L. interrogans* genogroup**

For genogroups G1-G13, significantly enriched KEGG pathways among genes with dN/dS >1 is listed, including gene counts and median dN/dS values. Fisher's exact test was used to estimate the enrichment of pathways. Significant pathways are highlighted in red.

**Supplementary Table 4. Genes under positive selection within SNP-enriched categories**
List of genes under positive selection (dN/dS >1) within SNP-enriched categories (cell wall/membrane, signal transduction, lipid A, peptidoglycan pathways) across genogroup. Genes under positive selection for each genogroup is indicated by a cross.

**Supplementary Table 5. Representative *L. interrogans* strains from genogroups G4-Ict and G13-Pom used in this study**

**Supplementary Table 6. Table of differentially expressed genes (*p*.adj ≤0.05) of *L. interrogans* G4-Ict in EMEM medium compared to EMJH medium at 3 days post incubation**
Specific G4-Ict genes do not present in G13-Pom are highlighting in yellow. For absence of annotation in the nomenclature *L. interrogans* serovar Copenhageni Fiocruz L1-130 strain, the nomenclature of *L. interrogans* serovar Lai 56601 strain was used.

**Supplementary Table 7. Differentially expressed genes (*p*.adj ≤0.05) of *L. interrogans* G13 in EMEM medium compared to EMJH medium at 3 days post incubation**
Specific G13-Pom genes do not present in G4-Ict are highlighting in yellow and the nomenclature of *L. interrogans* isolate 201801210 was used for the locus name (**Source Data Fig.5**). For absence of annotation in the nomenclature *L. interrogans* serovar Copenhageni Fiocruz L1-130 strain, the nomenclature of *L. interrogans* serovar Lai 56601 strain was used.

**Supplementary Table 8. Table of differentially expressed genes (*p*.adj ≤0.05) of *L. interrogans* G4-Ict compared to *L. interrogans* G13-Pom in EMEM at 3 days post incubation**

**Supplementary Table 9. Proteome of *L. interrogans* genogroups G4-Ict compared to G13-Pom in EMEM medium at 3 days post-incubation**

**Supplementary Table 10. Differentially expressed genes (*p*.adj ≤0.05) of human macrophages infected with *L. interrogans* G4-Ict compared to G13-Pom at 6 hr post-infection**

**Supplementary Table 11. Differentially expressed genes (*p*.adj ≤0.05) of murine macrophages infected with *L. interrogans* G4-Ict compared to G13-Pom at 6 hr post-infection**

**Supplementary Table 12. Primers used in this study**

### SOURCE DATA

**Source Data Fig.2. Protein sequences of *L. interrogans* isolate 201801210**

Fasta file of protein sequences from *L. interrogans* isolate 201801210, annotated using Prokka version 1.14.5.

### REFERENCES


1. Seemann, T. Prokka: rapid prokaryotic genome annotation. *Bioinformatics* **30**, 2068–2069 (2014).
2. Emms, D. M. & Kelly, S. OrthoFinder: phylogenetic orthology inference for comparative genomics. *Genome Biol.* **20**, 238 (2019).
3. Nieves, C. *et al.* Horizontal transfer of the rfb cluster in Leptospira is a genetic determinant of serovar identity. *Life Sci. Alliance* **6**, (2023).
4. Sorensen, M. *et al.* Rapid microbial identification and colistin resistance detection via MALDI-TOF MS using a novel on-target extraction of membrane lipids. *Sci. Rep.* **10**, 21536 (2020).
5. Pětrošová, H. *et al.* Lipid A structural diversity among members of the genus Leptospira. *Front. Microbiol.* **14**, 1181034 (2023).

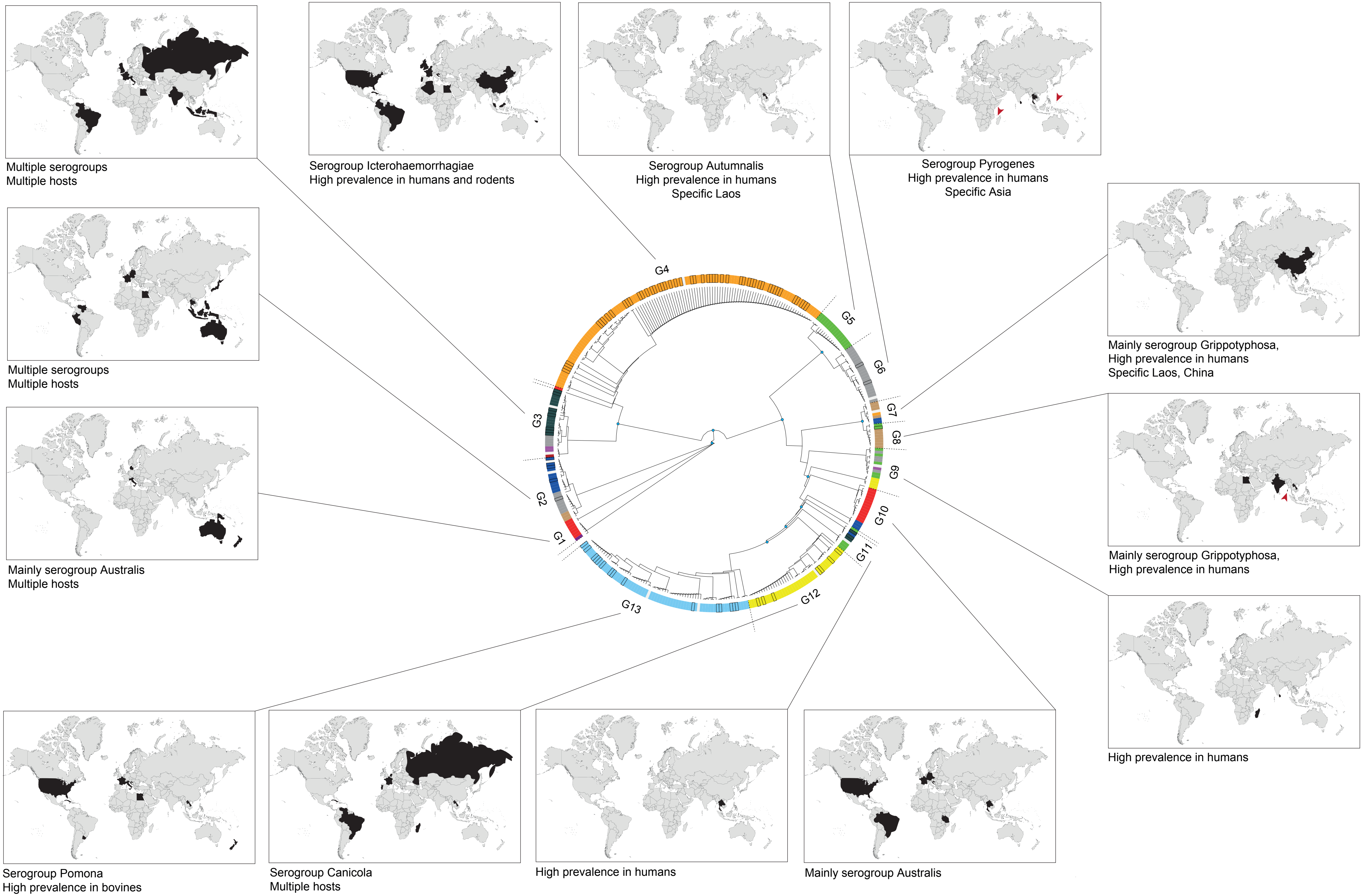


**Supplementary Fig. 1**

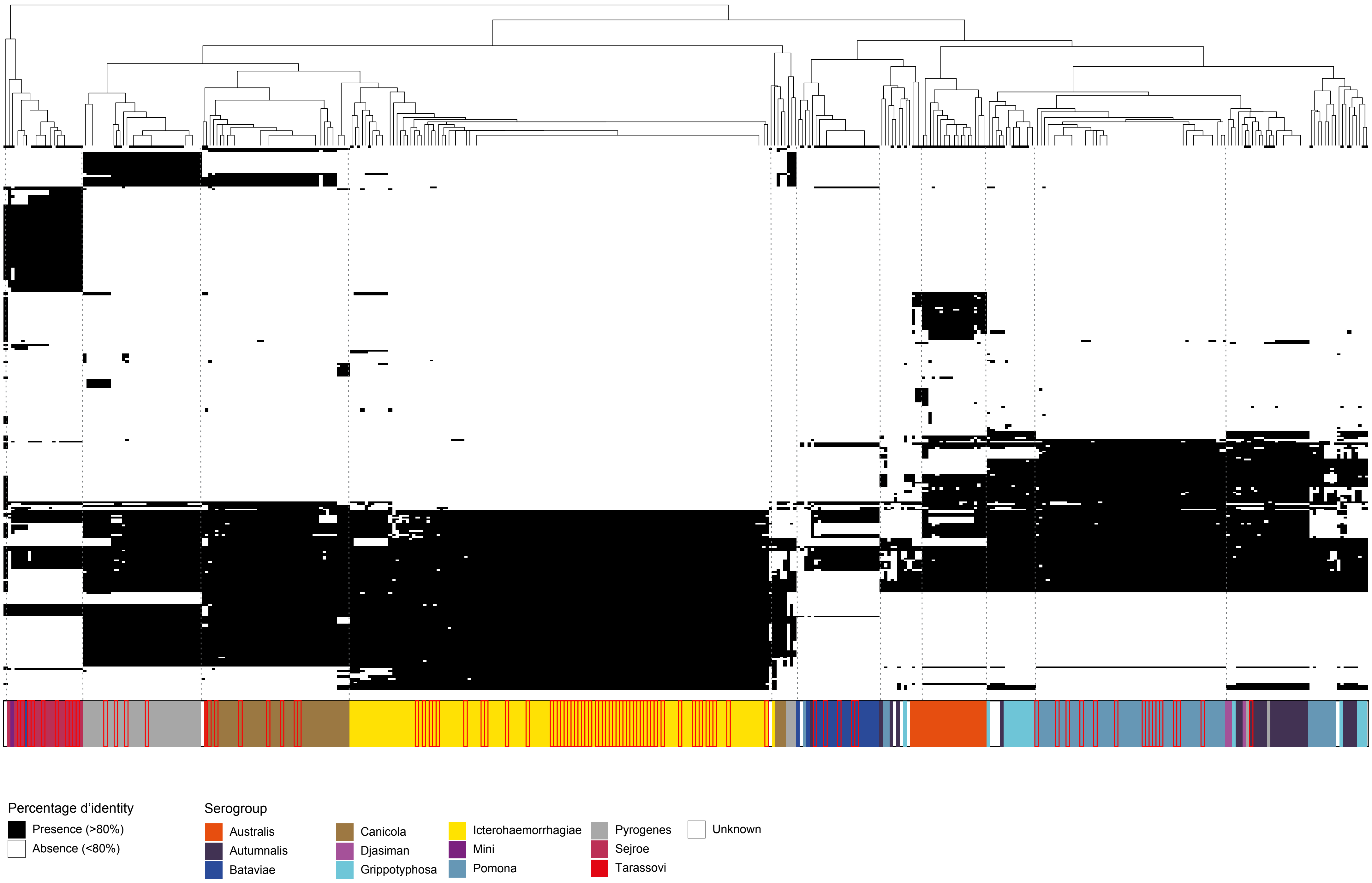


**Supplementary Fig. 2**

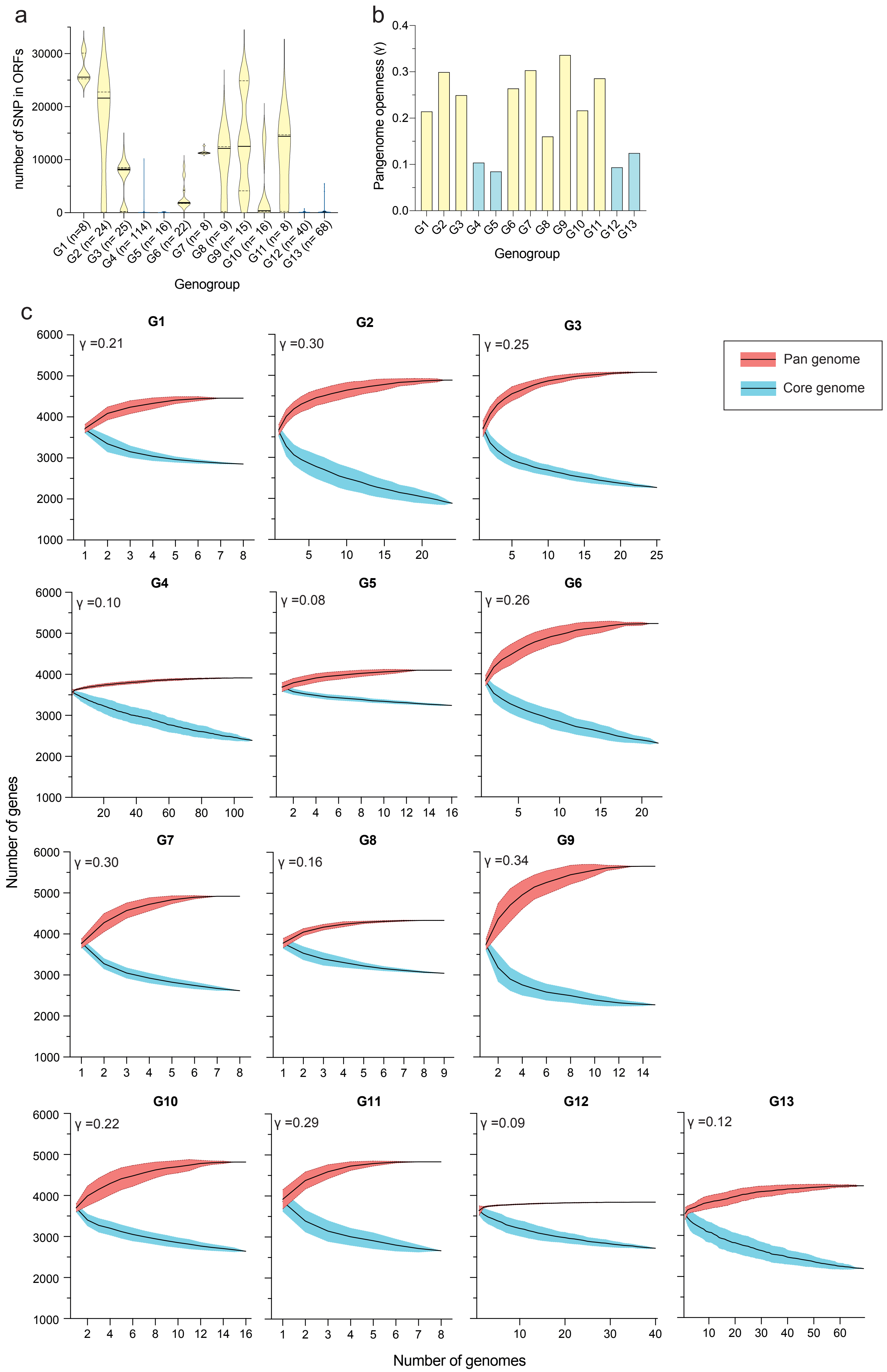


**Supplementary Fig. 3**

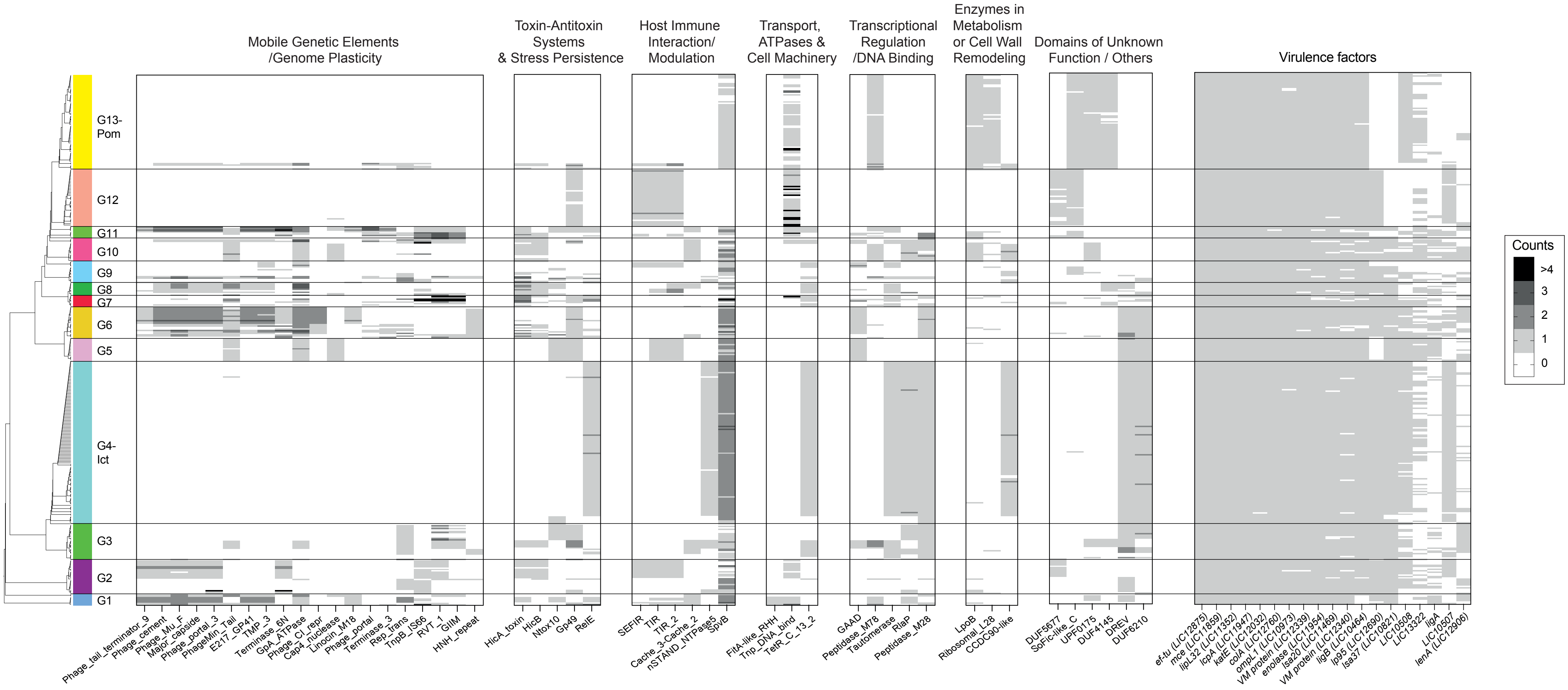


**Supplementary Fig. 4**

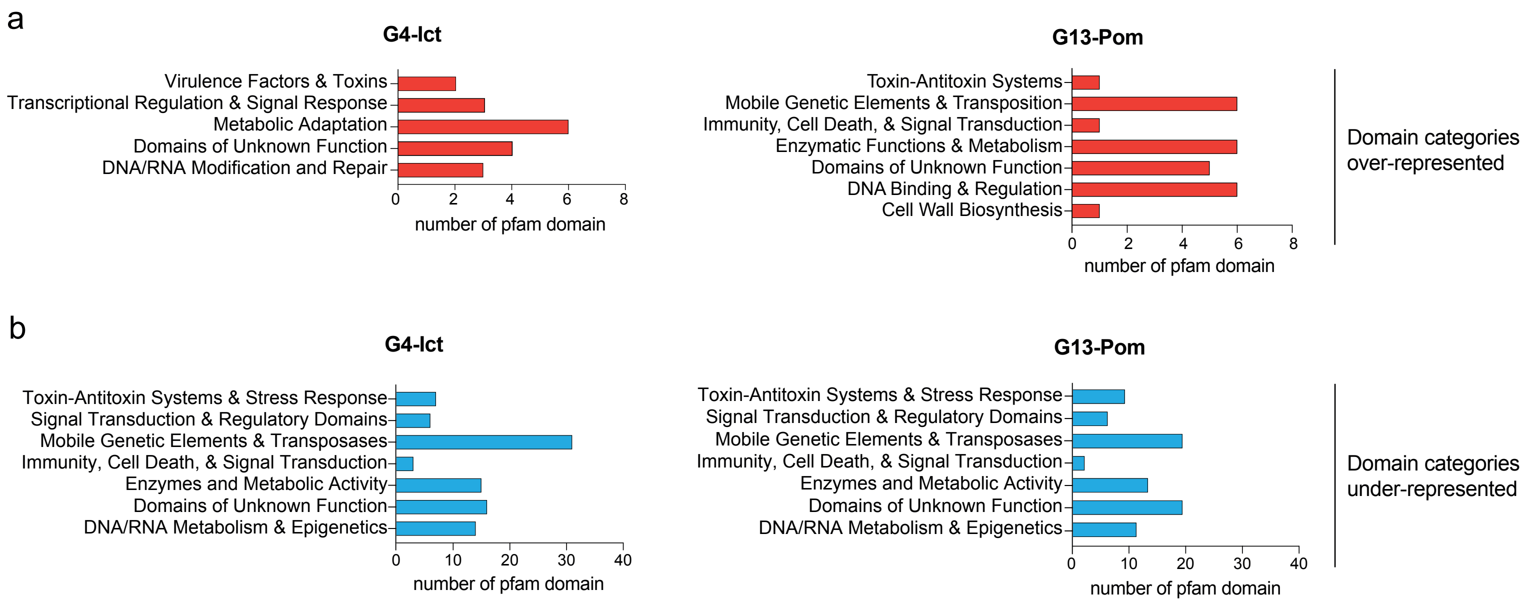


**Supplementary Fig. 5**

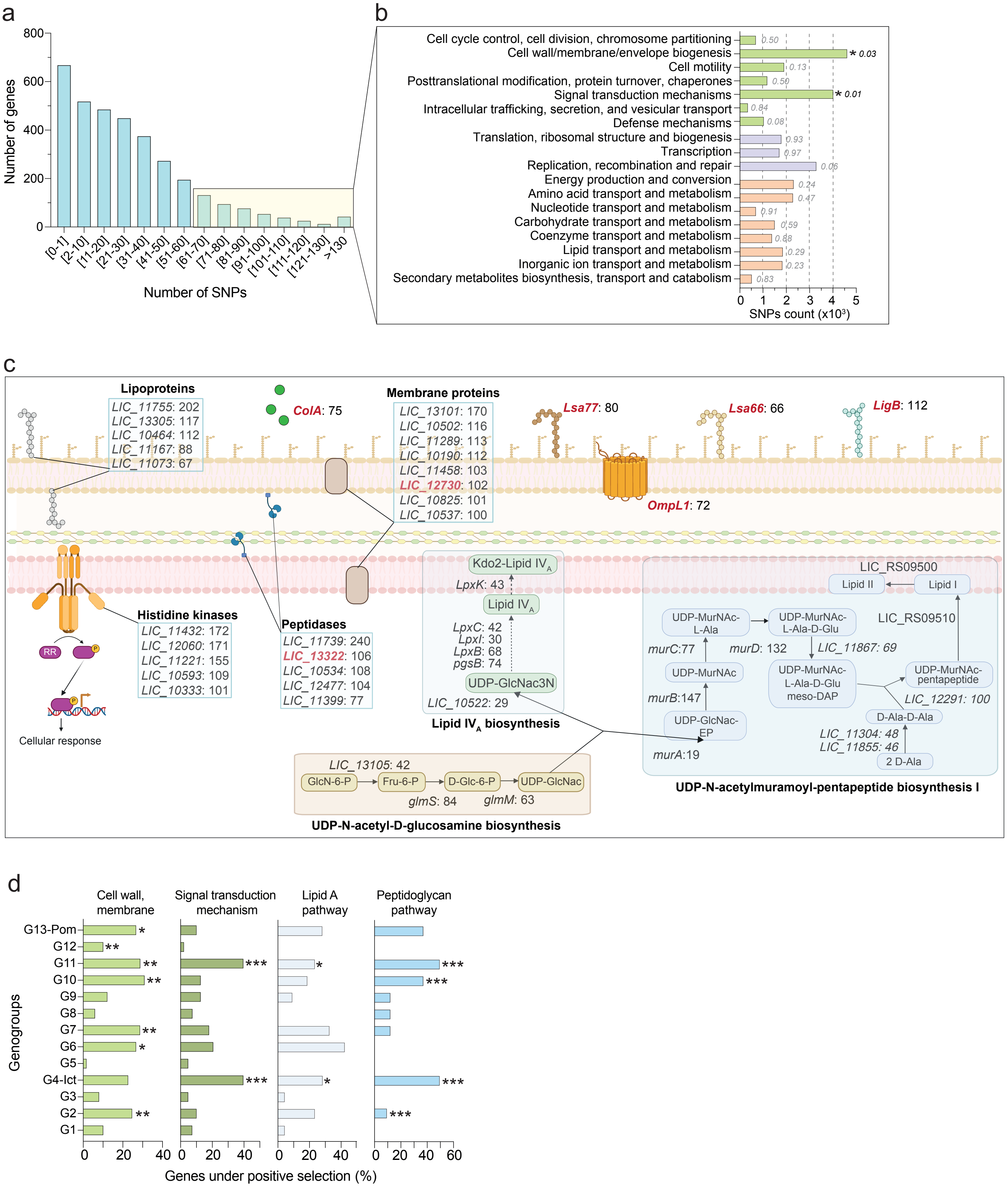


**Supplementary Fig. 6**

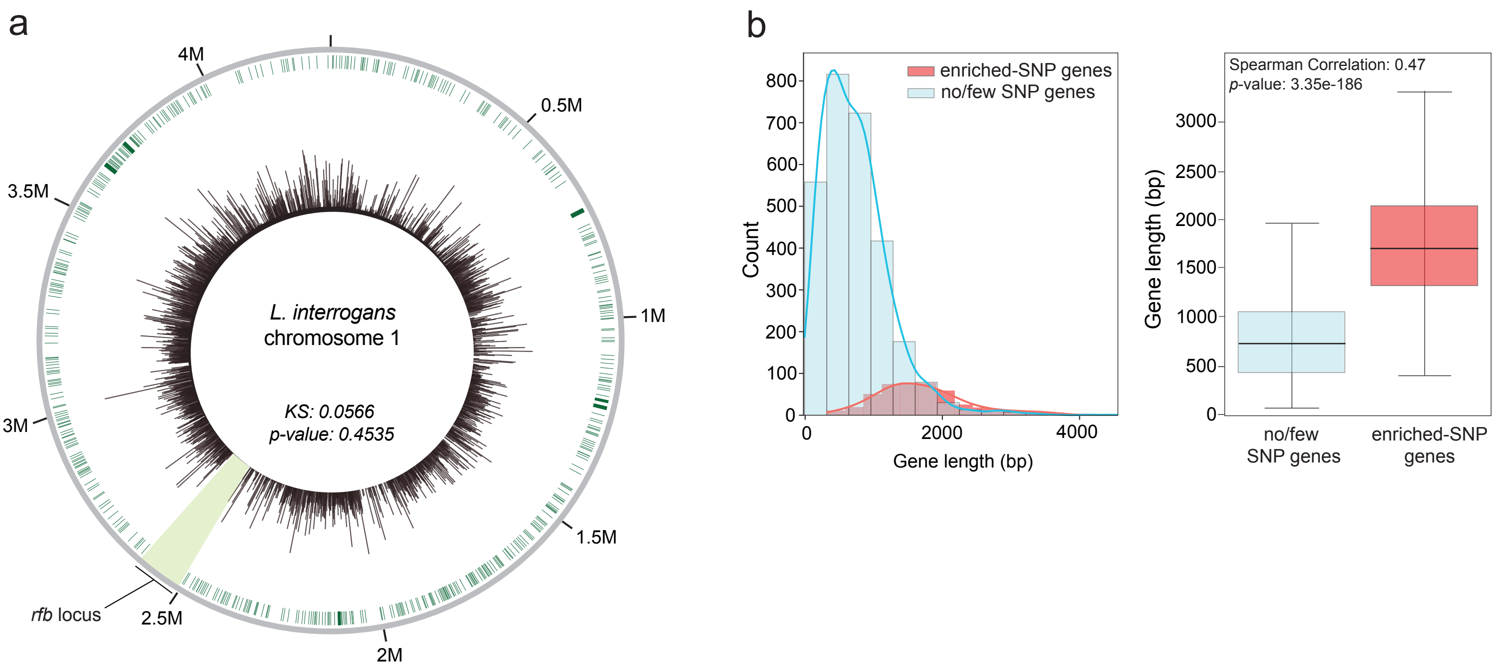


**Supplementary Fig. 7**

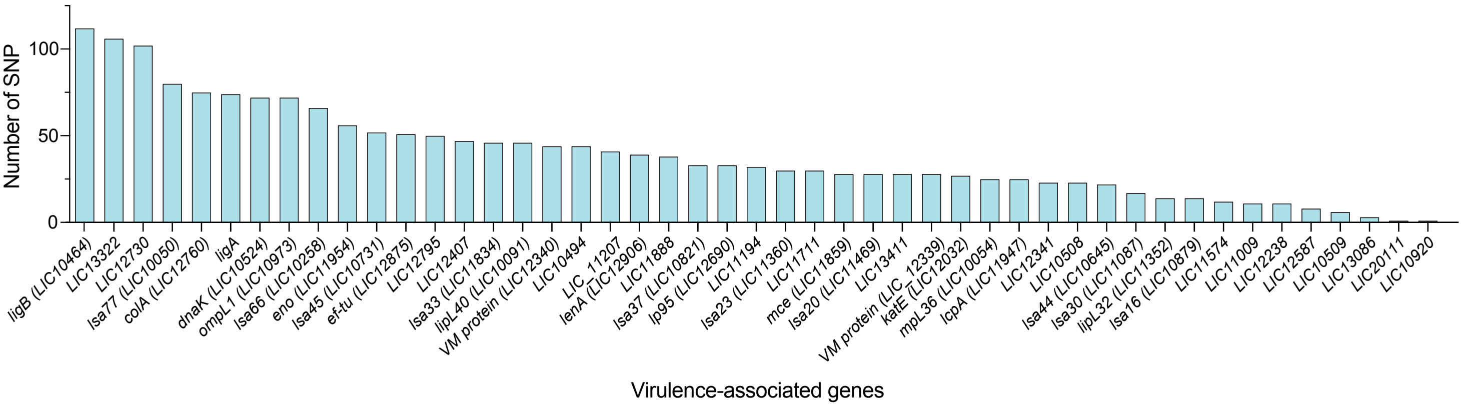


**Supplementary Fig. 8**

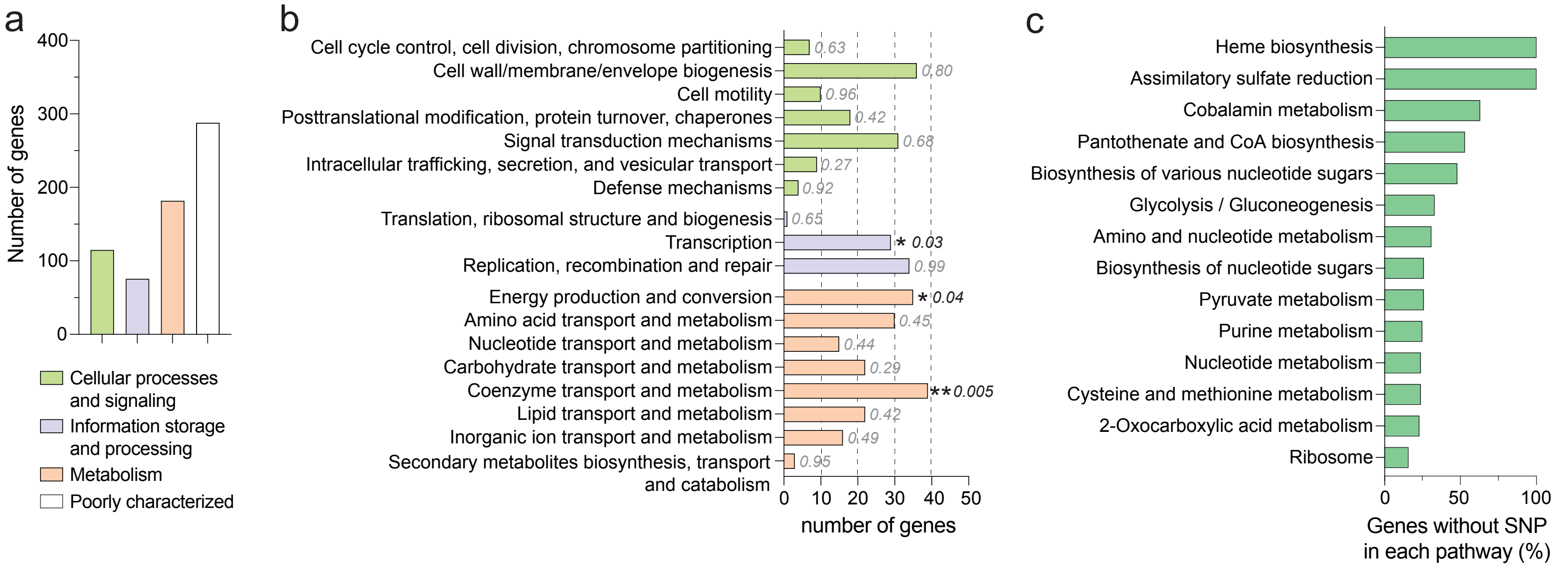


**Supplementary Fig. 9**

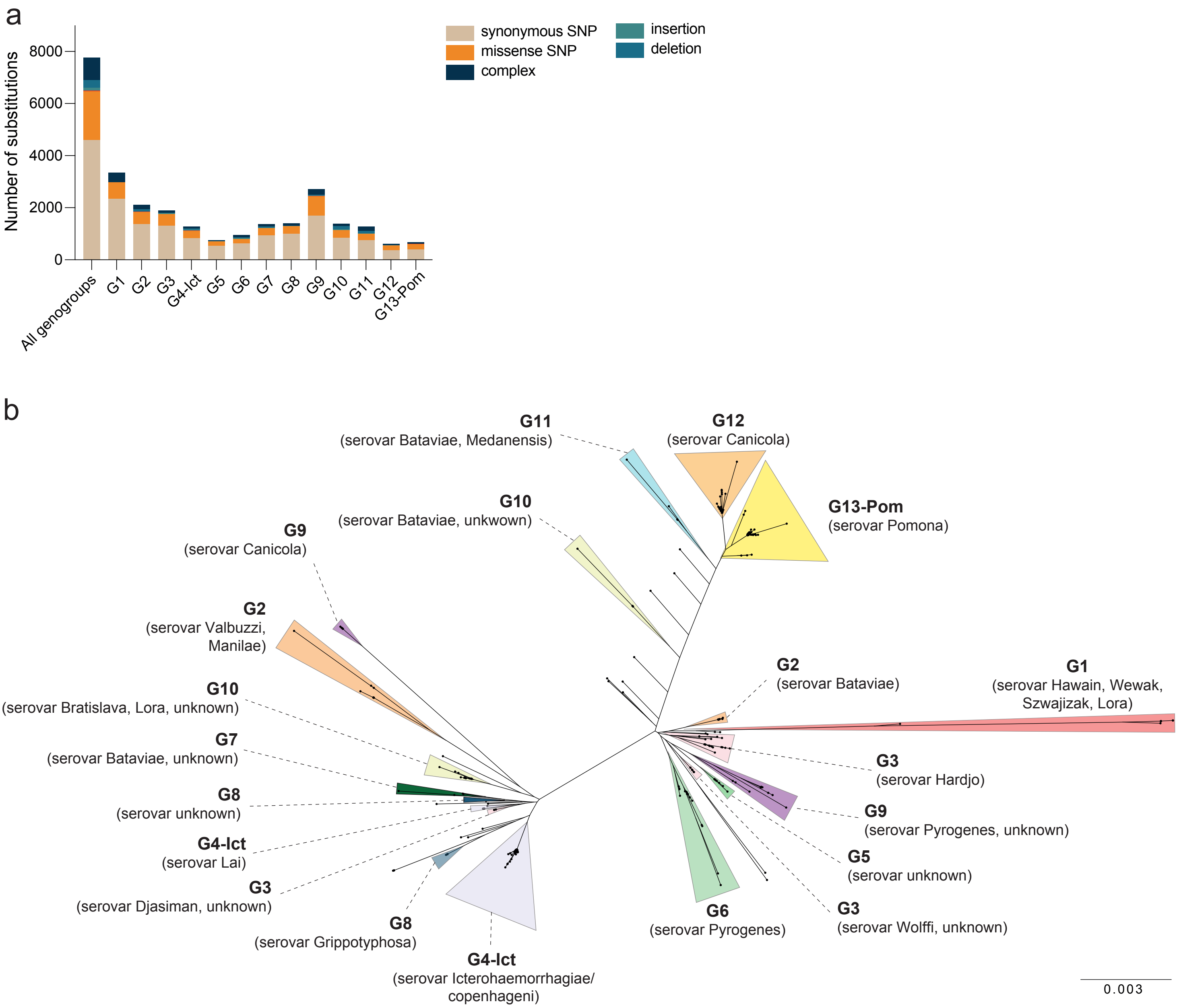


**Supplementary Fig. 10**

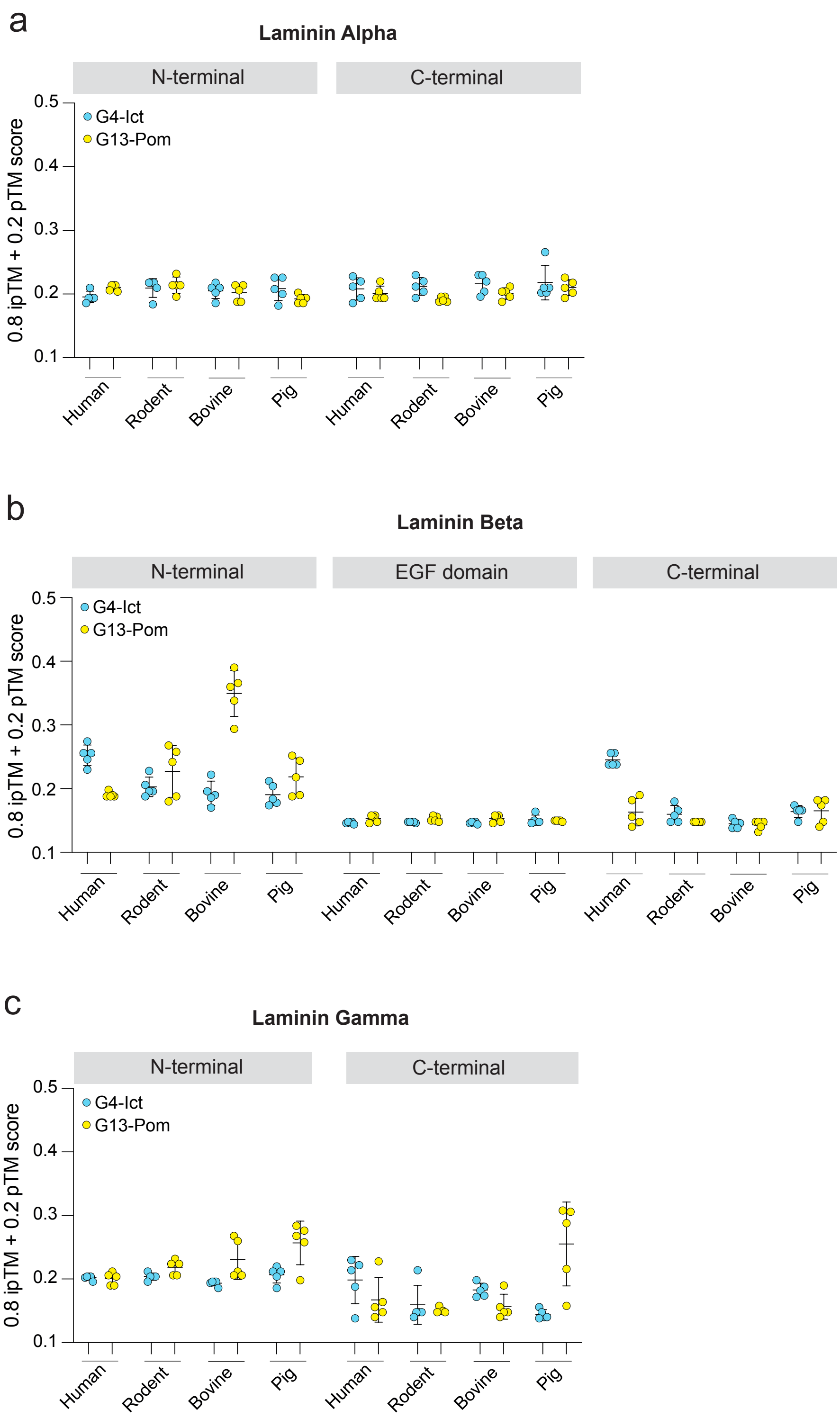


**Supplementary Fig. 11**

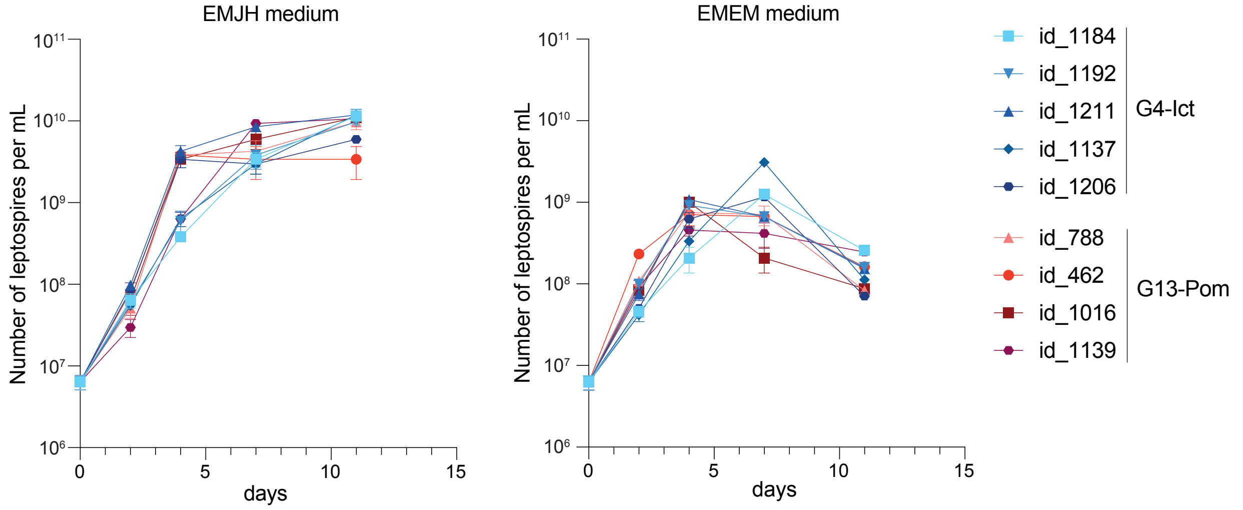


### Supplementary Fig. 12

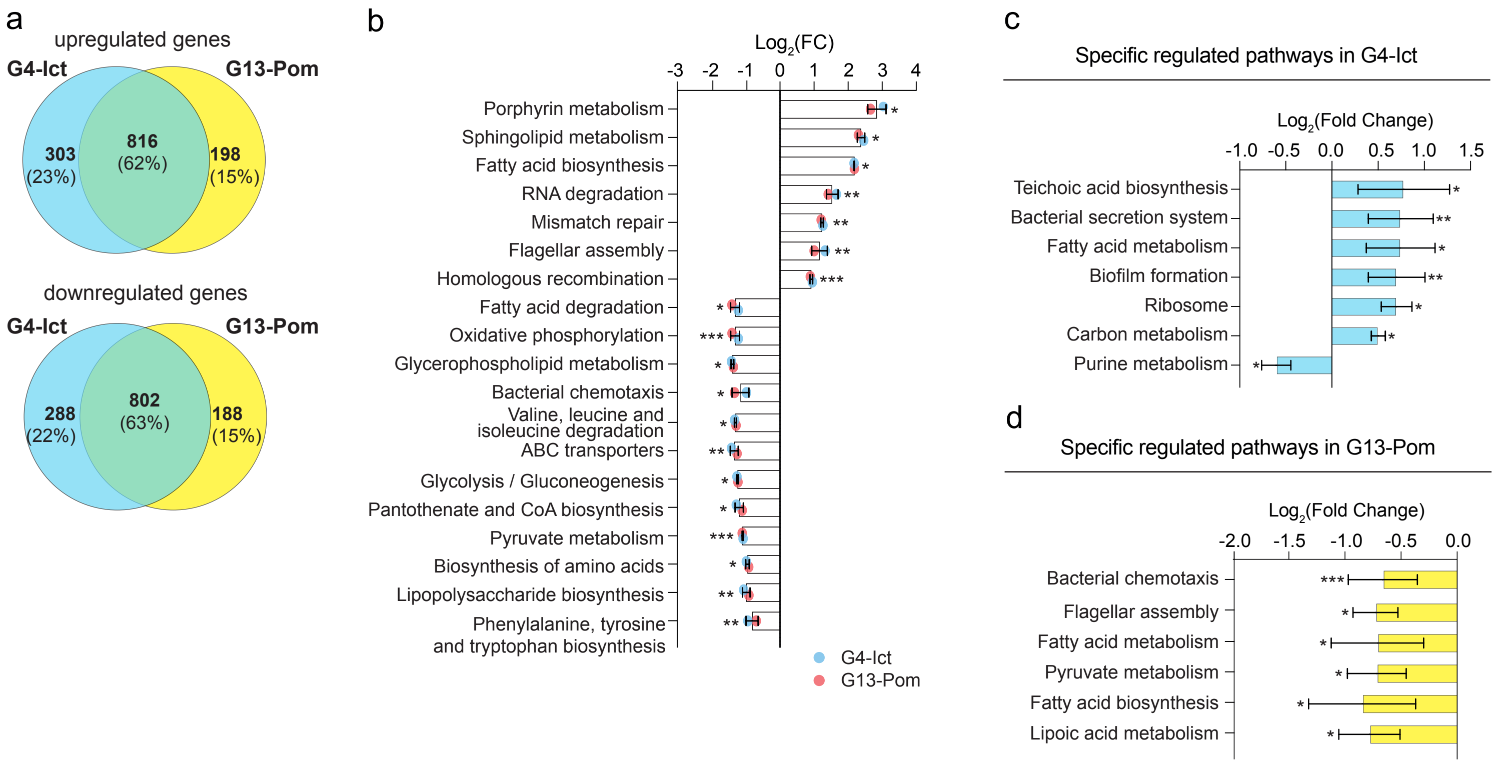


### Supplementary Fig. 13

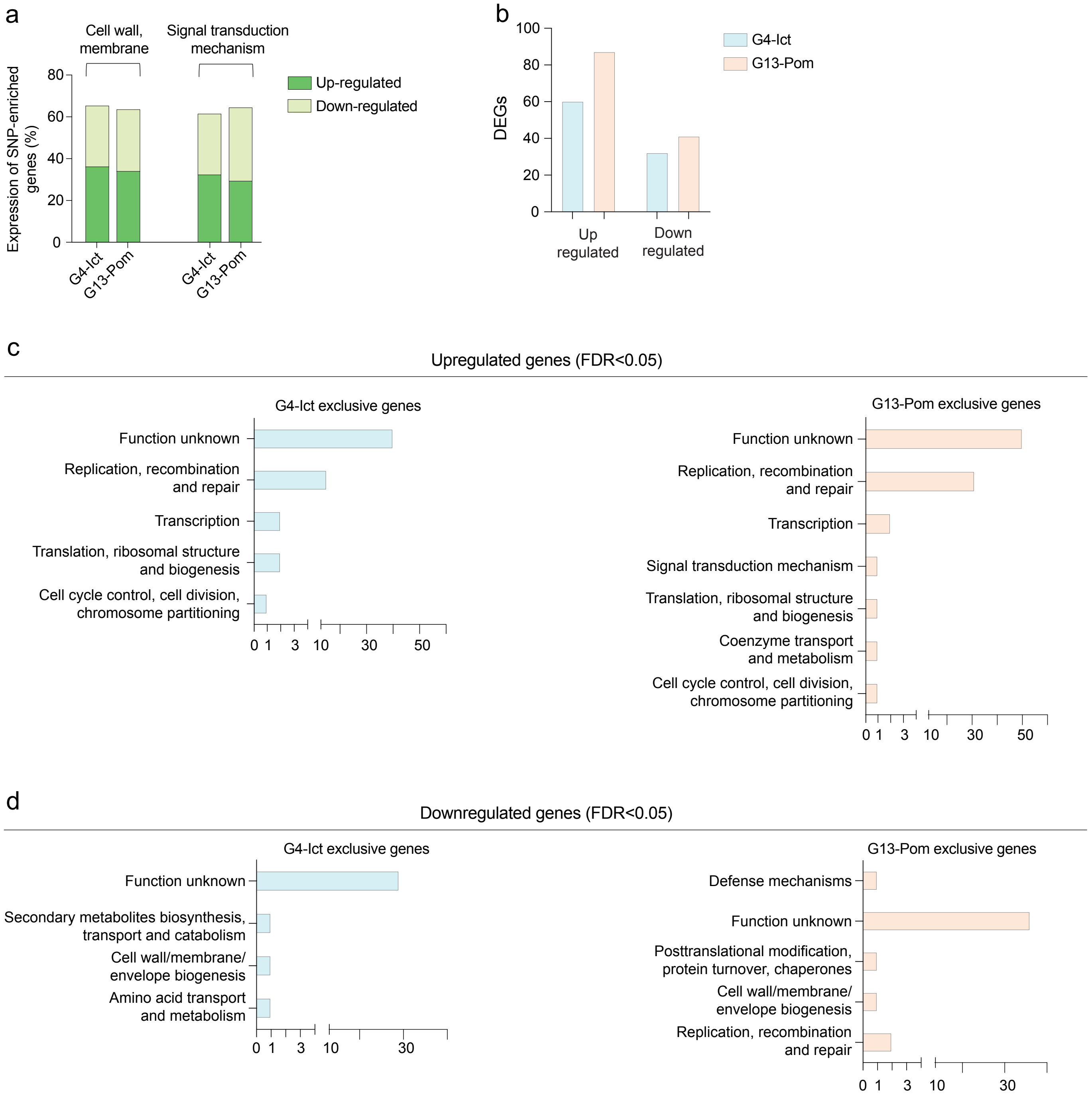


**Supplementary Fig. 14**

20 most-upregulated genes in G4-Ict vs G13-Pom

| gene name | $Log_2$(FC) |
|---|---|
| *LIC_11629 (dsbA)* | *2.83* |
| *LIC_12736 (lipoprotein)* | *2.53* |
| *LIC_11966 (ErpY-Like Lipoprotein)* | *2.06* |
| *LIC_10064* | *1.86* |
| *LIC_10373 (lipoprotein)* | *1.78* |
| *LIC_11502* | *1.71* |
| *LIC_11862* | *1.63* |
| *LIC_13416 (sodium transporter)* | *1.62* |
| *LIC_10011 (lpL21)* | *1.52* |
| *LIC_12099 (lipoprotein)* | *1.46* |
| *LIC_12340 (VM protein)* | *1.39* |
| *LIC_13037 (amino acid transporter)* | *1.38* |
| *LIC_11352 (Lipl32)* | *1.34* |
| *LIC_10094 (long-chain-fatty-acid CoA ligase)* | *1.34* |
| *LEPIC_0291* | *1.26* |
| *LIC_11729 (fadH)* | *1.24* |
| *LIC_11861* | *1.12* |
| *LIC_10464 (ligB)* | *1.05* |
| *LIC_10377 (lipoprotein)* | *0.99* |
| *LIC_11985 (RNA-binding protein)* | *0.97* |

20 most-upregulated genes in G13-Pom vs G4-Ict

| gene name | $Log_2$(FC) |
|---|---|
| *LIC_12675* | *3.58* |
| *LIC_10165 (nuclease inhibitor)* | *2.78* |
| *LIC_11687 (endonuclease)* | *2.60* |
| *LIC_12017 (chaperone protein ClpB)* | *2.42* |
| *LIC_10552 (histidine kinase)* | *2.40* |
| *LIC_10166* | *2.32* |
| *LIC_11426 (Phosphoserine phosphatase)* | *2.01* |
| *LIC_12158 (hydroxyacid aldolase)* | *1.60* |
| *LIC_12676* | *1.47* |
| *LIC_11425 (histidine kinase)* | *1.30* |
| *LIC_12627 (histidine kinase)* | *1.30* |
| *LIC_12183 (glycosyltransferase)* | *1.22* |
| *LIC_20172 (lipoprotein)* | *1.17* |
| *LIC_11788 (hydrolase)* | *1.06* |
| *LIC_20269 (alginate O-acetyltransferase)* | *1.04* |
| *LIC_10092 (metallo-beta-lactamase)* | *1.04* |
| *LIC_12188 (methyltransferase)* | *1.04* |
| *LIC_12210 (Hsp15-like protein)* | *1.02* |
| *LIC_12638 (Iron-sulfur cluster assembly scaffold protein IscU)* | *1.01* |
| *LIC_11263* | *1.00* |

### Supplementary Fig. 15

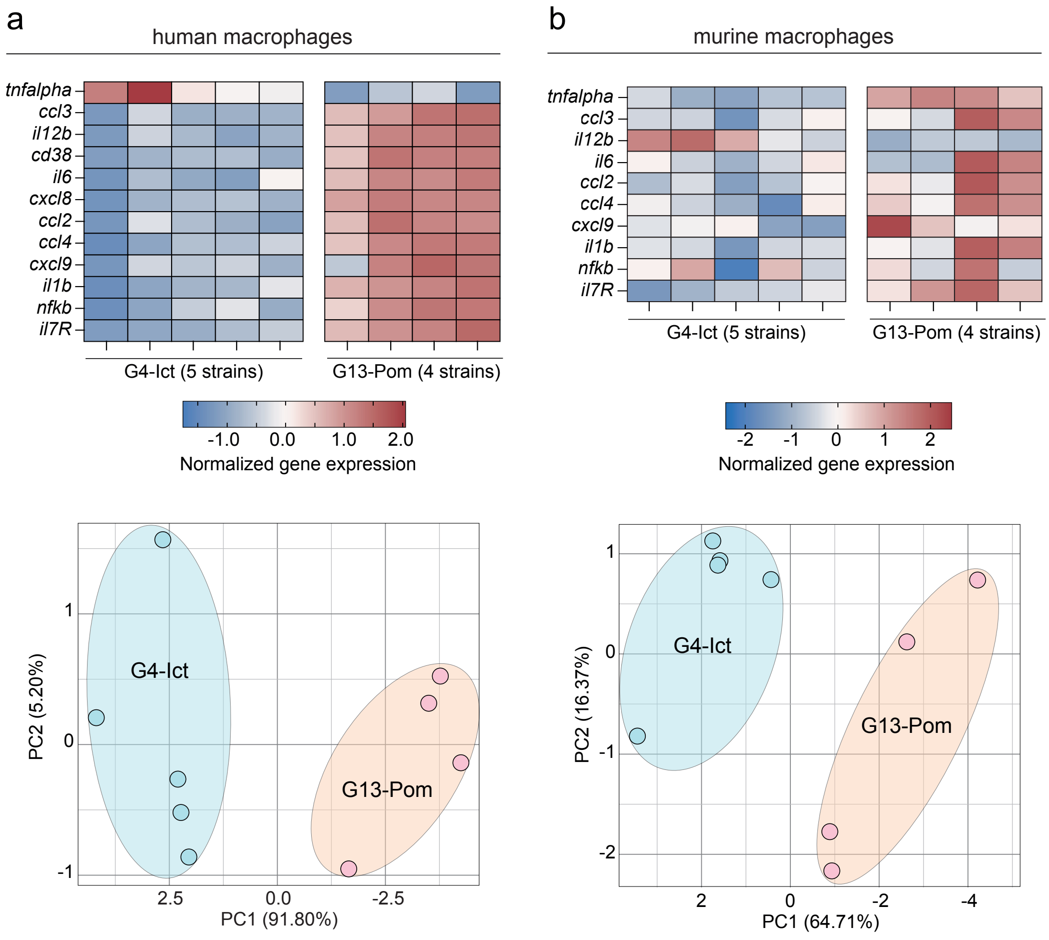


**Supplementary Fig. 16**

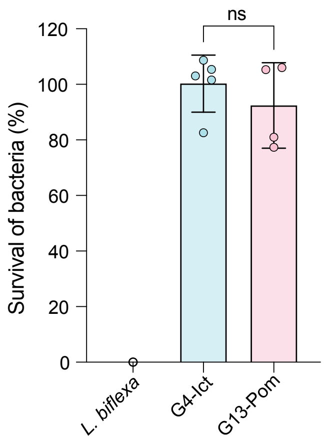


### Supplementary Fig. 17

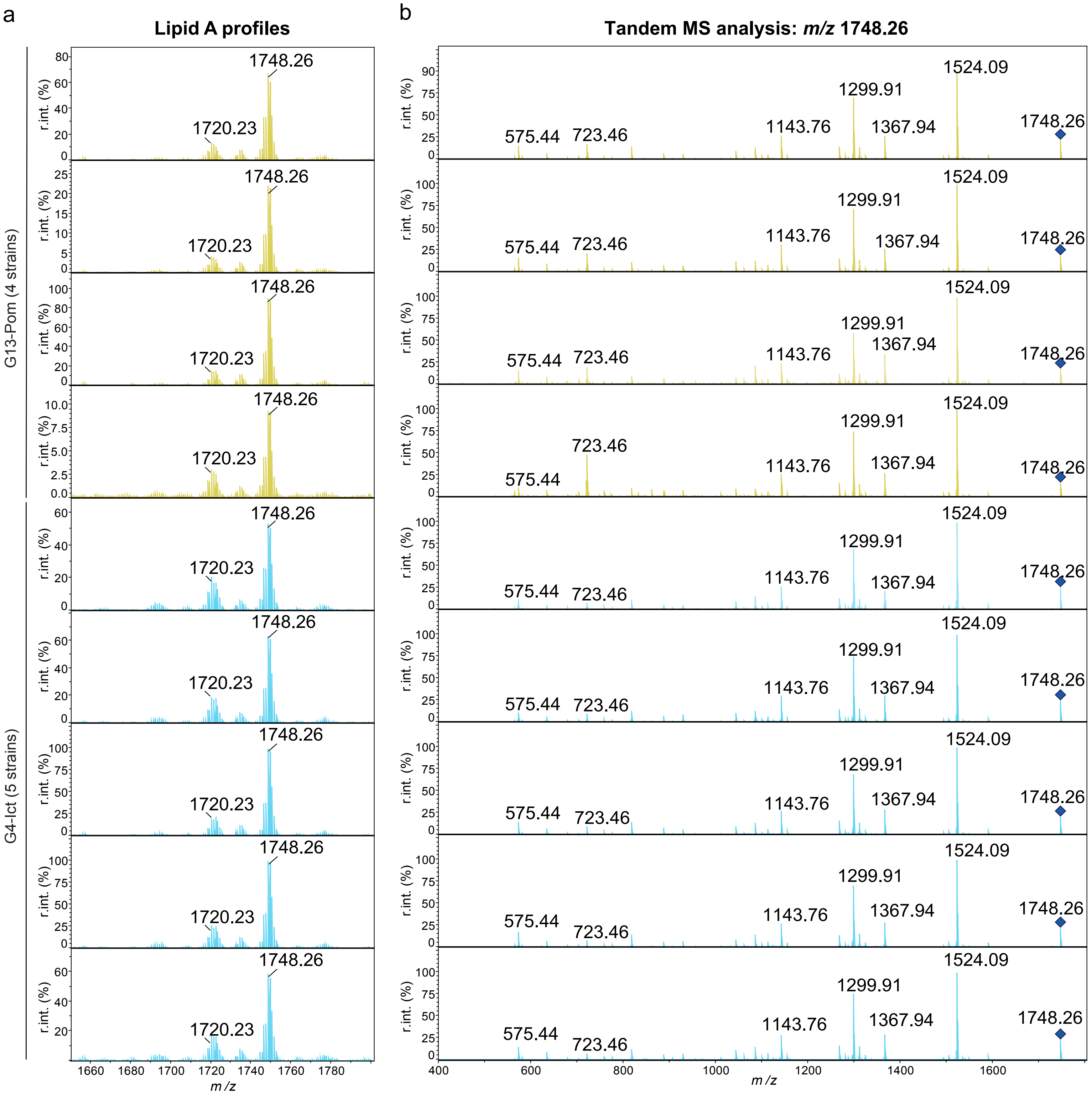


**Supplementary Fig. 18**

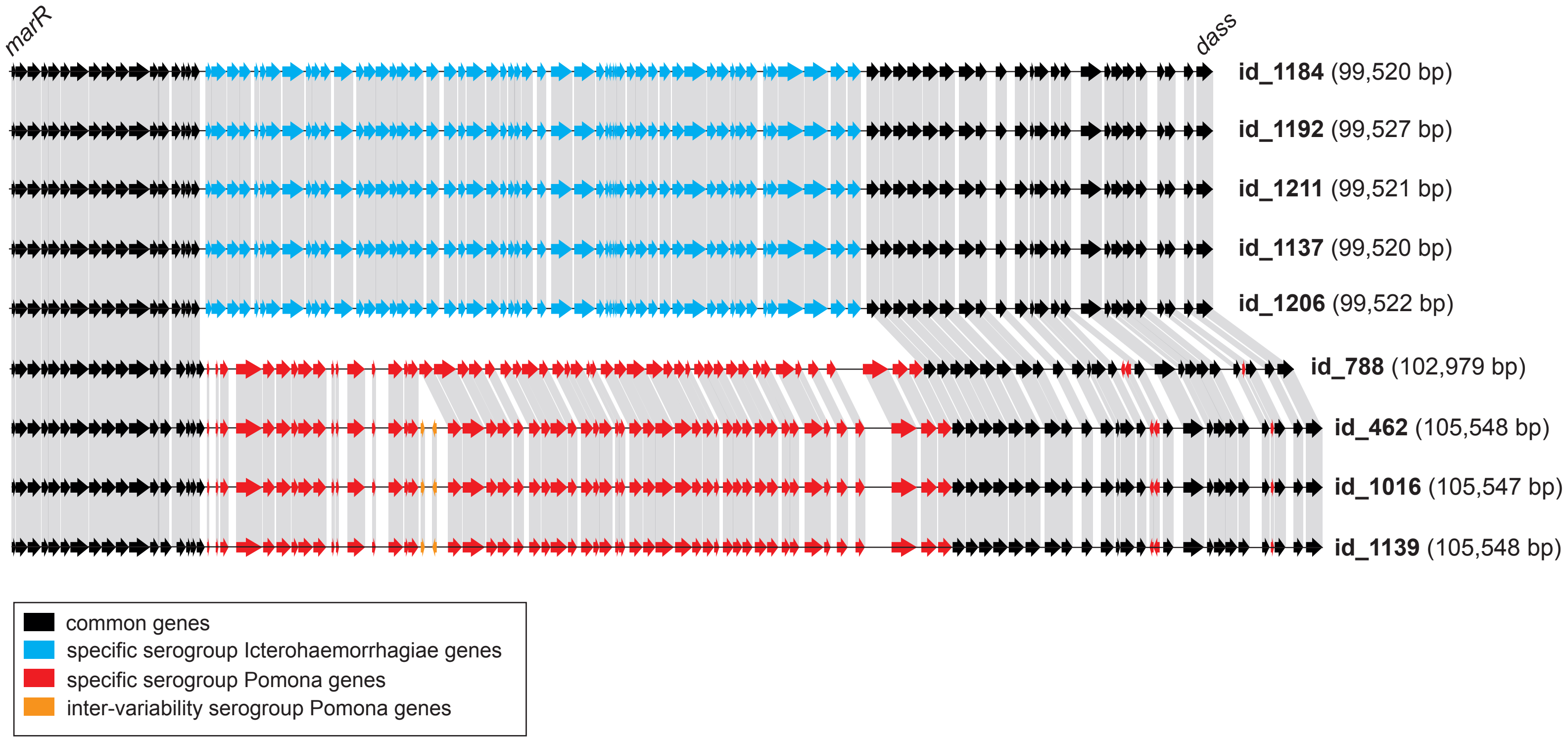


**Supplementary Fig. 19**
